# Depth-resolved analysis reveals target-dependent cryo-FIB damage profiles

**DOI:** 10.64898/2026.09.16.752186

**Authors:** Robert Michael Nwokonko, M. Zuhaib Qayyum, Dimple Karia, Vishal Maingi, Amrita Rai, Yuewen Sheng, Syed Asfarul Haque, Christopher Thompson, Sanja Sviben, Yanagisawa Haruaki, Jose Miguel De la Rosa-Trevin, Ravi Kalathur, Georgios Skiniotis, Abhay Kotecha

## Abstract

Cryogenic focused ion beam (cryo-FIB) milling enables cellular cryo-electron tomography but can compromise high-resolution structural information near milled surfaces. Here we establish recombinant human apoferritin (apoF) as a reporter of depth-dependent structural preservation in cellular lamellae. ApoF yielded a 2.05 Å *in situ* reconstruction. Apparent FIB damage was most pronounced within the first 15 nm for xenon, 30 nm for gallium, 45 nm for argon and remained detectable at least 75 nm from oxygen-milled surfaces under the workflows tested. Endogenous 70S ribosomes in the same xenon-milled tomograms showed a broader 30-45 nm profile, whereas β-galactosidase showed a similar narrow xenon-associated profile in a different specimen. Xenon-milled lamellae 125-150 nm thick yielded the highest resolution among the thickness groups examined, with lower resolution in the thinner and thickest groups. These findings show structural preservation is constrained by both surface-associated damage and lamella thickness, with the apparent extent of damage varying by molecular reporter. Previous ribosome-based surface-exclusion estimates may therefore be overly conservative for some molecular targets.

## Introduction

Cryo-electron tomography (cryo-ET) combined with subtomogram averaging (STA) enables high-resolution structure determination of macromolecular complexes within their native cellular environment while preserving molecular organization and intracellular interactions [1, 2]. Attainable resolution depends on specimen thickness and quality of generated lamella. As thickness increases, inelastic and multiple electron scattering reduce image contrast and effective signal-to-noise ratio and attenuate high-spatial-frequency information [3]. Because most cells and tissues are too thick for transmission electron microscopy, controlled thinning of vitrified specimens is often essential for high-resolution *in situ* analysis [3–6].

Cryogenic focused ion beam (cryo-FIB) milling has become central to preparing electron-transparent cellular lamellae for cryo-ET [7]. Conventional instruments typically use gallium liquid-metal ion sources, whereas plasma FIB systems use xenon, argon, nitrogen or oxygen ions at higher beam currents for faster material removal [8–10]. These developments have extended cryo-lamella preparation to thicker specimens ranging from cells to tissues [10–13]. However, ion milling can damage lamellae, resulting in loss of high-resolution information. Ion implantation and collision cascades perturb vitrified material beneath the milled surface [8], creating a trade-off between electron transparency and preservation of high-resolution information [14–16].

Earlier studies assessed milling damage using high-resolution cryo-ET, STA, or two-dimensional template matching (2DTM). For 30 keV gallium milling, reported apparent damage depths range from approximately 30-60 nm, with lower-energy polishing reducing the affected region [14–16]. Surface-associated information loss has also been detected after argon and xenon plasma FIB milling [8–10]. A recent ribosome-based comparison indicated better preservation with xenon than gallium or argon under the conditions tested [17]. Most measurements in biological lamellae have been performed on ribosomes, structurally robust ribonucleoprotein complexes approximately 25-30 nm in diameter [8–10, 14, 16, 17]. Because each ribosome spans a range of depths within the lamella, assigning its depth from its center limits how precisely surface-associated information loss can be localized.

Partial ablation, molecular composition, conformational heterogeneity, particle detection, and alignment accuracy may further influence the measured profile. Apparent damage depth therefore reflects both ion-specimen interactions and reporter properties and should not be interpreted as a fixed physical boundary.

Apoferritin (apoF), a conformationally homogeneous, octahedrally symmetric 24-subunit protein cage approximately 12 nm in diameter, is widely used to benchmark cryo-EM instrumentation and image-processing workflows [18, 19]. Its diameter is less than half that of the ribosome, reducing uncertainty in assigning particle depth from its center. Recombinant expression permits comparison after purification and within cells, while its symmetry and homogeneity enable robust reconstruction from small particle subsets required for depth-resolved analysis. The recent introduction of genetically encoded symmetric particles as intracellular standards for *in situ* cryo-ET further supports the use of defined molecular probes for quantitative benchmarking [20].

Here we expressed human apoF in *Escherichia coli* (*E. coli*) to quantify depth-dependent preservation of high-resolution structural information in cryo-FIB milled cellular lamellae. We determined purified and *in situ* apoF structures at near-atomic resolution and compared xenon, gallium, argon, and oxygen milling workflows. By analyzing endogenous 70S ribosomes within the same xenon-milled lamellae and extending the approach to recombinant β-galactosidase, we tested how reporter properties influence the estimation of apparent damage depth.

## Results

### Establishing a reference system with purified and *in situ* apoF

We prepared purified and *in situ* human apoferritin (apoF) specimens from the same *E. coli* expression culture. Some cells were plunge-frozen intact and thinned using a 30 kV xenon plasma ion beam; apoF from the remaining culture was purified and plunge-frozen on cryo-EM grids (Fig. 1a). This allowed comparison of apoF in thin vitreous ice and within a crowded cellular milieu.

**Figure 1.**
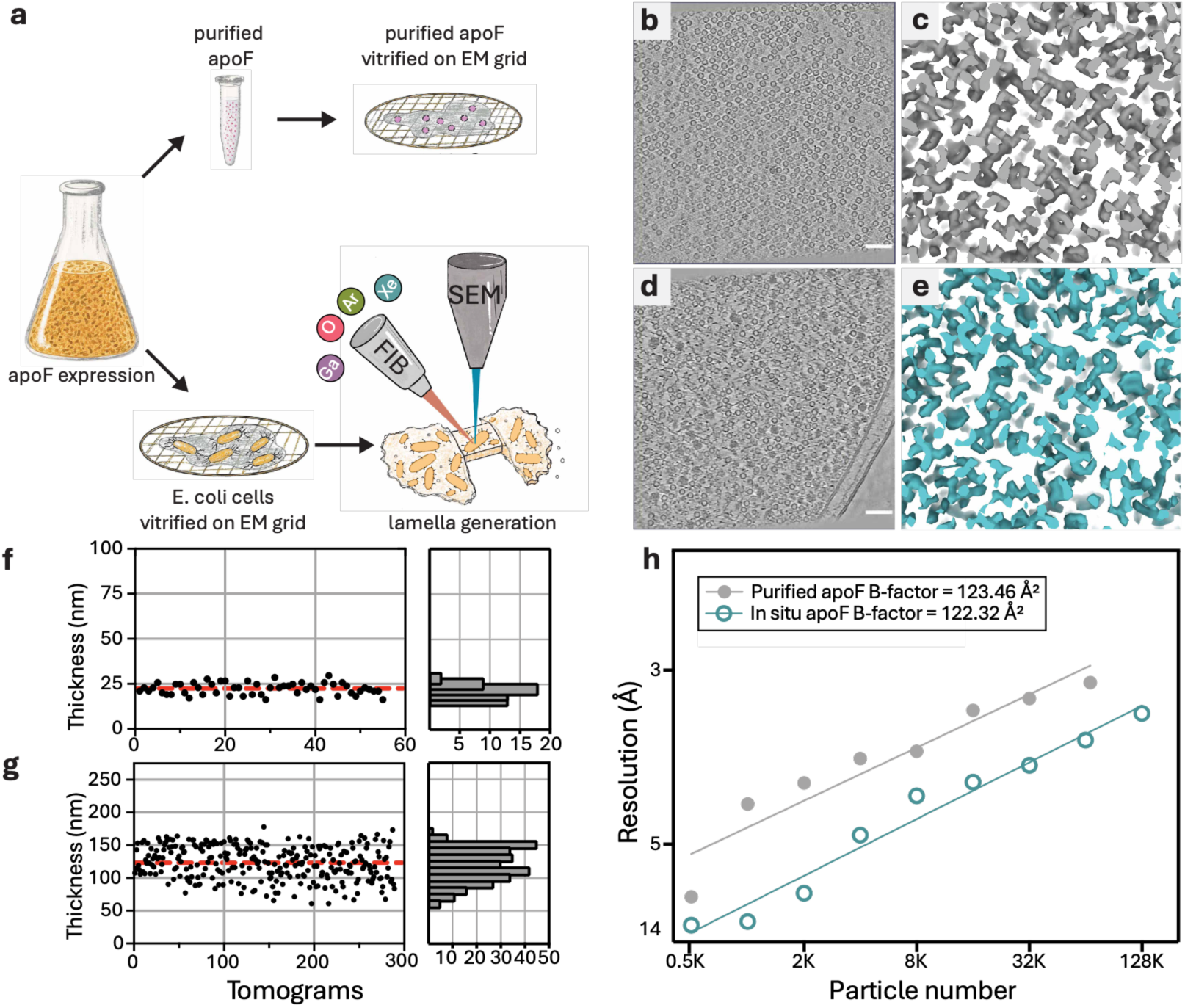
Sample preparation and high-resolution cryo-ET of purified and *in situ* apoferritin. a, Overview of apoferritin expression in *E. coli*, followed by either protein purification and vitrification or direct vitrification of cells and FIB milling. **b**, Denoised tomographic slice of purified apoferritin in vitreous ice. Scale bar, 50 nm. **c,** Density around a fourfold symmetry axis in the 1.97 Å reconstruction of purified apoferritin. **d,** Denoised tomographic slice of xenon FIB-milled *E. coli* lamella. Scale bar, 50 nm. **e,** Density around a fourfold symmetry axis in the 2.05 Å reconstruction of *in situ* apoferritin from xenon-milled lamellae. **f**, Ice thickness measurements from 55 tomograms of purified apoferritin (mean ± s.d., 22.4 ± 3.2 nm). **g**, Lamella thickness measurements from 289 tomograms of xenon-milled *E. coli* (mean ± s.d., 128.15 ± 25.4 nm). Red dashed lines in **f** and **g** represents the estimated mean thickness. **h,** Rosenthal-Henderson analysis of purified (gray) and *in situ* (cyan) apoferritin, yielding apparent B-factors of 123.5 and 122.3 Å², respectively.

We acquired 55 tilt series from the purified specimen and 447 from xenon-milled cellular lamellae (Supplementary Table 1). After reconstruction, we manually excluded tomograms with poor alignment, freezing or ice artifacts, apoF nanocrystals or no detectable apoF particles. We retained all 55 purified apoF and 289 cellular tomograms, with particles readily identifiable in thin vitreous ice and the bacterial cytoplasm, respectively (Fig. 1b, d).

Purified apoF was embedded in vitreous ice with a mean thickness of 22.4 ± 3.2 nm, whereas xenon-milled lamellae had a mean thickness of 128 ± 25.4 nm (Fig. 1f, g). Cellular apoF was therefore imaged in approximately fivefold thicker specimens, within the compositionally complex bacterial cytoplasm and between two ion-beam-exposed surfaces.

Tilt series were acquired at a physical pixel size of 1.2 Å, corresponding to a physical Nyquist limit of 2.4 Å. At physical-pixel (4K) sampling, Fourier shell correlation (FSC) curves for both reconstructions reached this limit. Data recorded in electron-event representation (EER) format with a Falcon 4i detector retain electron-event positions on a 16K super-resolution grid [21]. Rendering at 6K super-resolution sampling enabled refinement beyond the physical Nyquist limit, yielding global resolutions of 1.97 Å for purified apoF and 2.05 Å for *in situ* apoF (Supplementary Fig. 1). Both maps resolved the expected high-resolution features, including central holes in aromatic side chains (Fig. 1c, e). Rosenthal-Henderson analysis [22] yielded similar B-factors of 123.5 Å² for purified apoF and 122.3 Å² for *in situ* apoF (Fig. 1h), indicating comparable rates of resolution improvement with increasing particle number. For the same particle number, purified apoF reached somewhat higher resolution, consistent with a reduced signal-to-noise ratio in thicker, more compositionally complex cellular environments.

### Milling workflows differ in lamella thickness and information recovery

We next used apoF to compare structural preservation across four cryo-FIB workflows. All plasma FIB lamellae were initially milled with xenon to approximately 600 nm, then thinned and polished with xenon, argon or oxygen (Supplementary Fig. 2). We compared these with lamellae prepared using a gallium liquid-metal ion source (Supplementary Fig. 6). Plasma FIB final thinning used the same sequence of nominal aperture indices. However, equivalent aperture settings do not necessarily produce comparable beam currents, current densities, or probe profiles across ion species and instruments. These experiments compare complete milling workflows under the conditions tested rather than intrinsic effects of individual ion species.

The xenon workflow produced the thinnest lamellae, with a mean thickness of 128 ± 25.4 nm, compared to 171.0 ± 34.3 nm for argon, 178.0 ± 21.0 nm for gallium, and 209.0 ± 25.7 nm for oxygen (Fig. 2a). Further thinning of the argon, gallium, or oxygen-milled specimens compromised lamella integrity, preventing these workflows from reproducibly reaching the thicknesses achieved with xenon.

**Figure 2.**
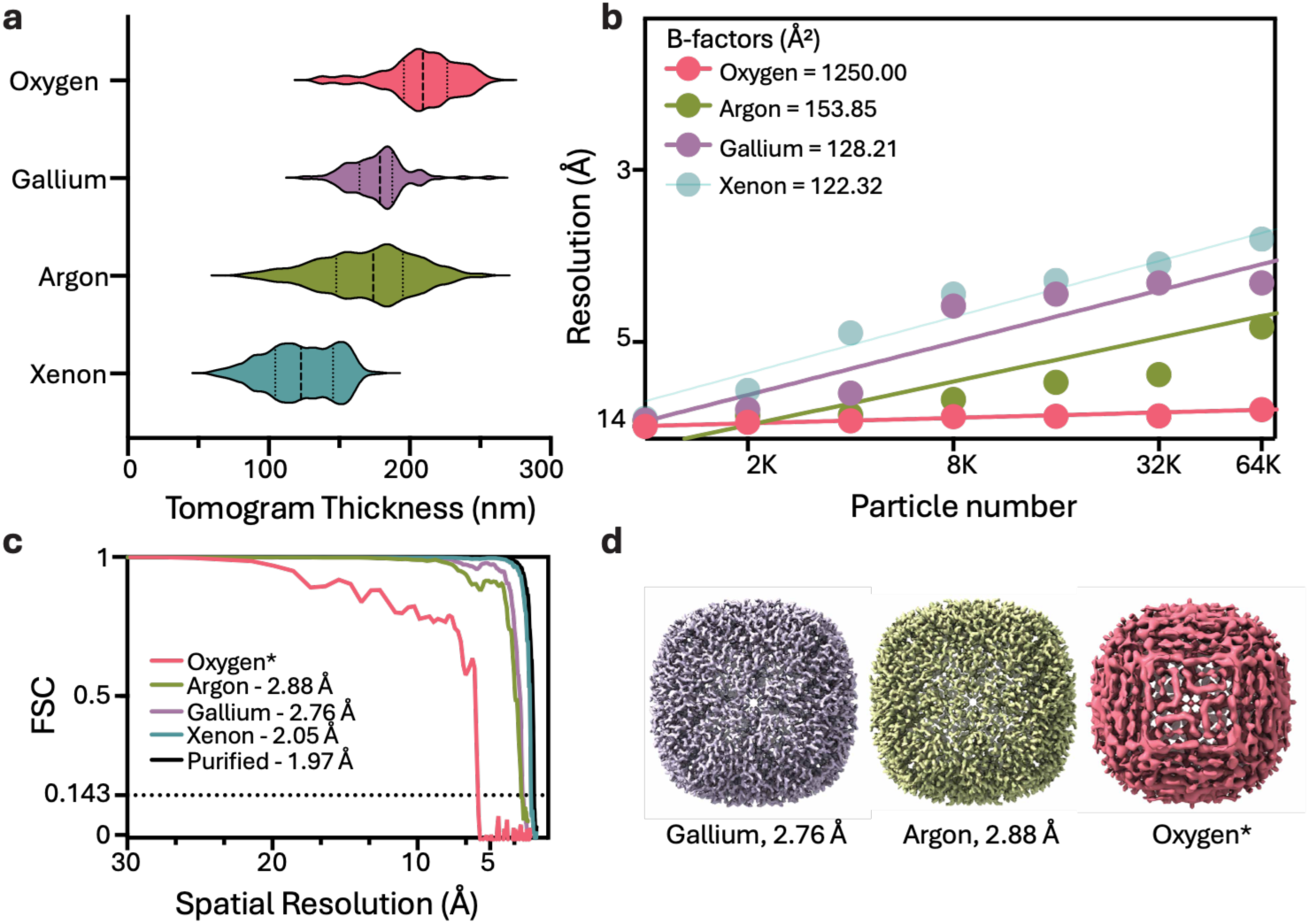
Comparing cryo-FIB milling workflows using different ion species. a, Distributions of estimated lamella thickness for xenon (128.15 ± 25.4 nm), argon (171.0 ± 34.3 nm), oxygen (209.0 ± 25.7 nm) and gallium milling (178.0 ± 21.0 nm). **b,** Rosenthal– Henderson analysis of *in situ* apoferritin, yielding B-factors of 1,250 Å² for oxygen, 153.85 Å² for argon, 128.21 Å² for gallium and 122.32 Å² for xenon. **c,** Fourier shell correlation (FSC) curves for purified apoferritin (1.97 Å) and *in situ* apoferritin from xenon (2.05 Å), gallium (2.76 Å), argon (2.88 Å) and oxygen (*) datasets. **d,** Apoferritin reconstructions from lamellae milled with gallium (purple), argon (green) and oxygen (red). *The oxygen reconstruction did not reproduce the expected apoF subunit arrangement, consistent with unreliable particle alignment; its FSC-derived resolution was therefore not reported.

Rosenthal-Henderson analysis yielded B-factors of 122.3 Å² for xenon, 128.2 Å² for gallium, and 153.9 Å² for argon (Fig. 2b). The oxygen dataset yielded a B-factor of approximately 1,250 Å² and little resolution improvement as particle number increased. Mean lamella thickness alone did not explain these differences: gallium and argon-milled lamellae had similar mean thicknesses, yet gallium yielded a lower B-factor.

Gold-standard FSC analysis yielded global resolutions of 2.05 Å for xenon, 2.76 Å for gallium and 2.88 Å for argon (Fig. 2c). The gallium and argon maps reproduced the characteristic apoF cage, whereas the oxygen map failed to reproduce the expected subunit arrangement, suggesting particle misalignment (Fig. 2d). The nominal FSC estimate for oxygen was therefore not considered a reliable measure of structural resolution, and the reconstruction is marked with an asterisk rather than a resolution assignment.

Excluding apoF particles within 75 nm of the nearest oxygen-milled surface improved particle alignments and yielded a 3.60 Å reconstruction. B-factors also decreased from 1,250 Å² to 465 Å² despite fewer particles (Supplementary Fig. 4; Supplementary Fig. 8). Although these results do not establish that particles beyond 75 nm were physically undamaged, they show that inclusion of surface-proximal particles led to poor resolution in the oxygen dataset.

### Lamella thickness affects reconstruction resolution within the xenon dataset

To assess the relationship between lamella thickness and high-resolution information, we grouped xenon-milled tomograms by measured lamella thickness into 60-100, 100-125, 125-150 and 150-175 nm and evaluated resolution and B-factors (Supplementary Fig. 3). B-factors decreased from approximately 111 Å² for 60-100 nm lamellae to 103 Å² for 125-150 nm lamellae, before increasing again to approximately 106 Å² for 150-175 nm lamellae. Reconstruction resolution followed similar trend, with the highest resolution obtained from 125-150 nm thick lamellae.

We also evaluated MultiShot acquisition in Tomography 5 software to increase acquisition throughput and assess the potential effects of beam-image shift induced aberrations on reconstruction quality. Using beam-image shifts of up to 3 µm, MultiShot enabled collection from two to three positions per acquisition and yielded a 2.31 Å *in situ* apoF reconstruction. The single-position and MultiShot datasets had similar mean lamella thicknesses, but the MultiShot dataset contained a larger proportion of thick tomograms. B-factors were 122.3 Å² for single-position acquisition and 173.8 Å² for MultiShot. Comparing with similar lamella thickness yielded similar B-factors of 148.3 and 142.0 Å², respectively (Supplementary Fig. 10). These results supported the use of MultiShot for high-resolution tomography at the beam-image shifts tested.

### Surface-associated information loss varies across milling workflows

To assess depth-dependent damage, we used IMOD [23] to annotate the upper and lower milled surfaces in each tomogram and interpolated them to generate boundary models. For each apoF particle, we calculated the distance from its center to the nearest milled surface and grouped particles into 15 nm intervals from 0 to 75 nm, with a final bin spanning 75-105 nm (Fig. 3a).

**Figure 3.**
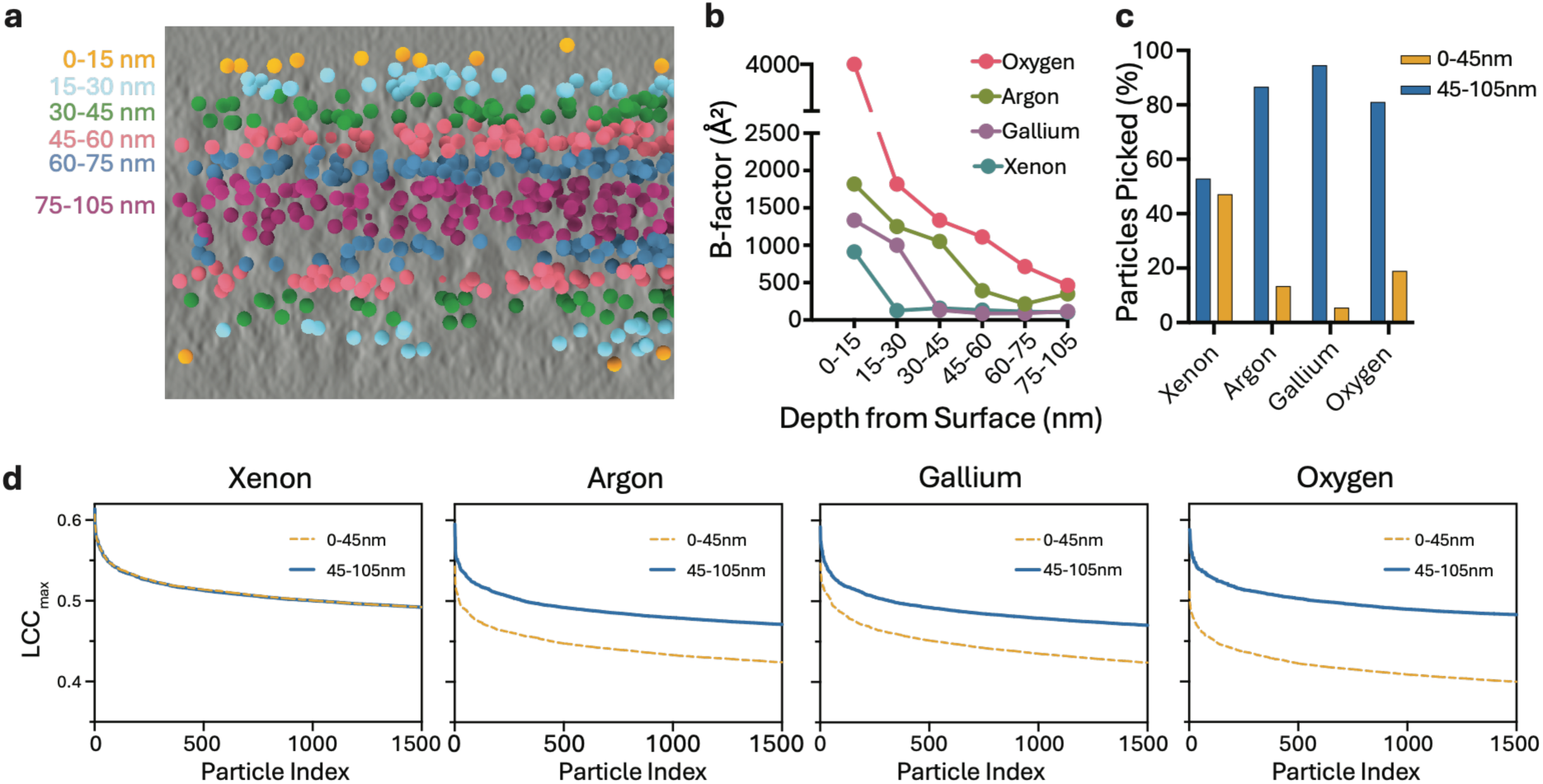
FIB induced damage profiles measured using apoferritin. a, Representative X– Z tomographic section showing apoferritin particle positions colored by depth, defined as the distance from each particle center to the nearest milled surface. **b,** B-factors estimated by Rosenthal–Henderson analysis of apoferritin reconstructions from successive depth intervals in xenon, argon, gallium and oxygen-milled lamellae. **c,** Percentages of detected apoferritin particles in the outer 0-45 nm (yellow) and the inner 45-105 nm (blue) depth groups for xenon, argon, gallium and oxygen milling workflows. **d,** Maximum locally normalized cross-correlation (LCC_max) scores for the 1500 highest scoring particles picked from each depth group: 0-45 nm (yellow) and 45-105 nm (blue). Scores are ranked from highest to lowest for xenon, argon, gallium and oxygen datasets.

Rosenthal-Henderson analysis showed different depth-dependent profiles across the four milling workflows (Fig. 3b). In the xenon dataset, the B-factor decreased sharply between the 0-15 nm and 15-30 nm bins and remained comparatively stable at greater depths (Supplementary Fig. 5a). In gallium-milled lamellae, B-factors remained elevated to approximately 30 nm before stabilizing (Supplementary Fig. 6), whereas in argon-milled lamellae B-factors remained elevated to approximately 45 nm (Supplementary Fig. 7). Oxygen produced the broadest profile, with markedly elevated B-factors throughout the first 75 nm and the lowest value only in the 75-105 nm interval; no clear plateau was observed within the sampled range (Supplementary Fig. 8). These apparent damage depths are operational estimates based on the depth-dependent loss of high-resolution information in apoF under the conditions tested, rather than discrete physical boundaries.

We also examined the depth distribution of apoF particles detected by template matching (Fig. 3c). In xenon-milled lamellae, 47% of particles were within 0-45 nm of the nearest surface and 53% in the deeper 45-105 nm region. Only 6% and 14% of detected particles in the gallium and argon datasets, respectively, were within the first 45 nm. The oxygen dataset contained a larger surface-proximal fraction, at 19%, although these particles yielded poor quality reconstruction and high B-factors.

To assess whether template-matching scores depended on depth, we compared LCC_max distributions for the 1,500 highest-scoring particles from each of the 0-45 nm and 45-105 nm ranges for each workflow (Fig. 3d). LCC_max is the maximum locally normalized cross-correlation between the target and template [24, 25]; higher values indicate stronger agreement with the apoF template. The xenon distributions showed little separation between depths. By contrast, selected particles within 45 nm of gallium, argon, or oxygen-milled surfaces had lower LCC_max values than particles from deeper regions. Oxygen yielded the largest delta between LCC_max values in both ranges (Fig. 3d).

### Apparent damage depth depends on the molecular reporter

The apparent damage profile measured using apoF in xenon-milled lamellae is narrower than previously reported ribosome-based estimates of 30-45 nm [8–10, 14, 16]. We therefore asked whether the measured xenon-associated profile depended on the molecular reporter. To compare reporters under the same experimental conditions, we identified endogenous 70S ribosomes in the xenon-milled tomograms used for apoF analysis and processed them independently by STA. The 70S ribosome and 50S subunit reconstructions reached global resolutions of 3.26 Å and 3.60 Å, respectively, with local resolution extending beyond 3 Å in both maps (Fig. 4b, c; Supplementary Fig. 11a, b).

**Figure 4.**
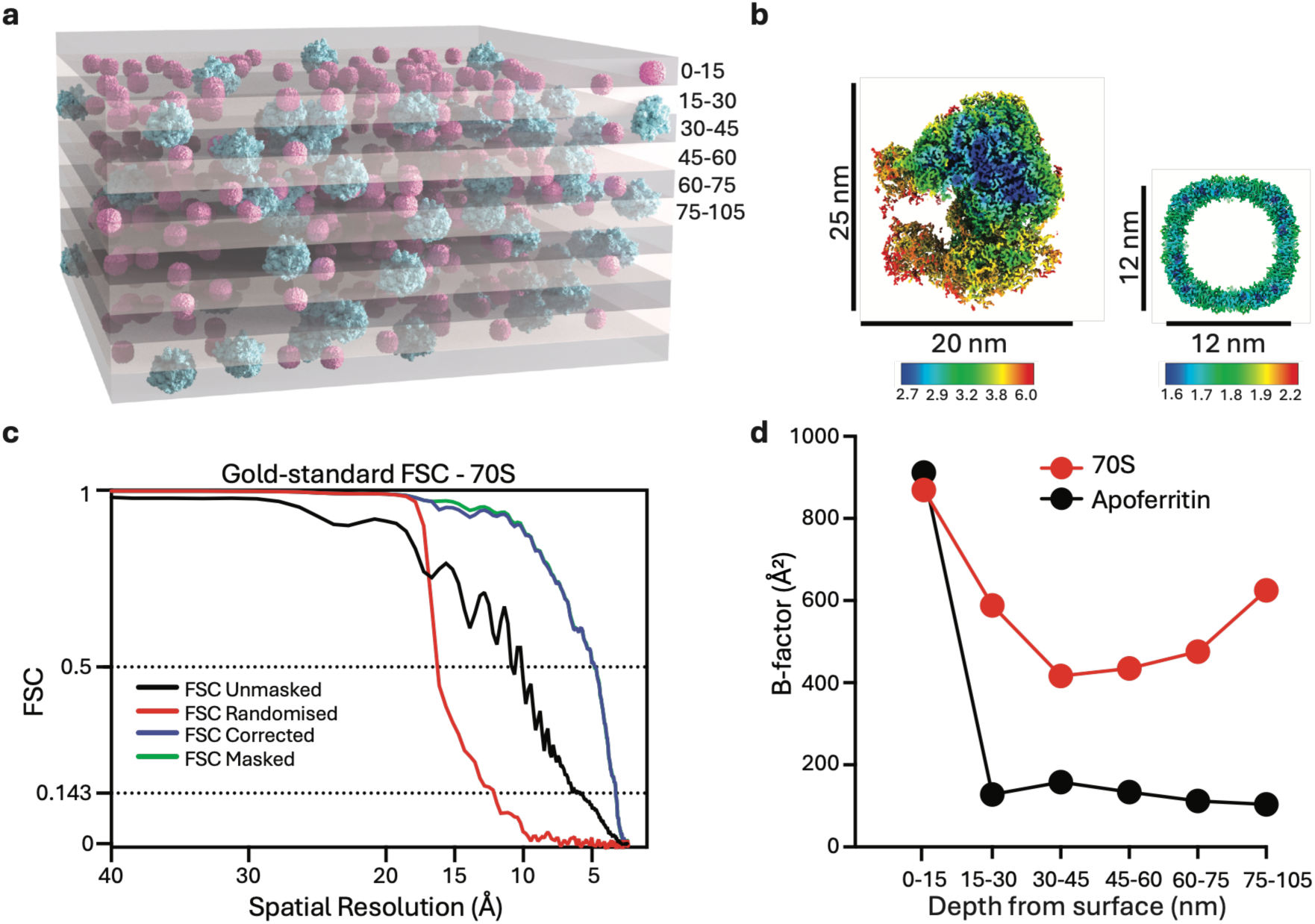
Comparison of cryo-FIB damage profiles measured using apoferritin and 70S ribosomes from the same tomograms. a, Three-dimensional representation of apoferritin (pink) and 70S ribosome (cyan) particles within a xenon-milled lamella, illustrating their assignment to depth intervals relative to the nearest milled surface. **b,** Size comparison of the 70S ribosome (approximately 25 × 20 nm) and apoferritin (approximately 12 nm in diameter). Reconstructions are colored by local resolution (Å) **c,** Gold-standard Fourier shell correlation (FSC) curves for the 70S ribosome reconstruction, indicating a resolution of 3.26 Å at FSC = 0.143 with local resolution extending beyond 3 Å as shown in **b**. **d,** B-factors estimated by Rosenthal–Henderson analysis for 70S ribosomes (red), plotted alongside previously determined apoferritin values (black) from the same xenon-milled tomograms. Particle depth was measured from each particle center to the nearest milled surface. Particles were grouped into 15 nm intervals from 0 to 75 nm, followed by a final 75-105 nm interval.

We then analyzed the 70S ribosomes using the same depth intervals previously used for apoF. Figure 4a, b illustrates these layers and the size difference between the reporters. Rosenthal-Henderson analysis showed ribosome B-factors remained elevated throughout the first 30 nm from xenon-milled surfaces and decreased substantially in the 30-45 nm interval, consistent with previously reported measurements [8–10, 14, 16]. For direct comparison, we plotted these values alongside the previously determined apoF B-factors, which decreased sharply beyond the first 15 nm and remained relatively stable at greater depths (Fig. 4d; Supplementary Fig. 5a, b). Because both reporters came from the same tomograms, this difference cannot be attributed to milling or specimen preparation and shows that apparent damage depth also depends on molecular target.

### β-galactosidase reproduces the narrow xenon damage profile

We next tested whether another protein could reproduce the narrow xenon-associated profile we saw for apoF. We selected *E. coli* β-galactosidase, a D2-symmetric homotetramer with a more extended multidomain structure, greater conformational flexibility, and lower symmetry than apoF. β-galactosidase is also a benchmark for high-resolution cryo-EM, with structures reported at 3.2 Å, 2.2 Å, and 1.8 Å [26–29].

*E. coli* overexpressing β-galactosidase were plunge-frozen and thinned using a 30 kV xenon plasma ion beam. STA yielded an *in situ* β-galactosidase reconstruction at a global resolution of 2.94 Å (Fig. 5a, c, d). The β-galactosidase and apoF lamellae had similar thickness distributions, allowing comparison of their depth-dependent profiles (Fig. 5b).

**Figure 5.**
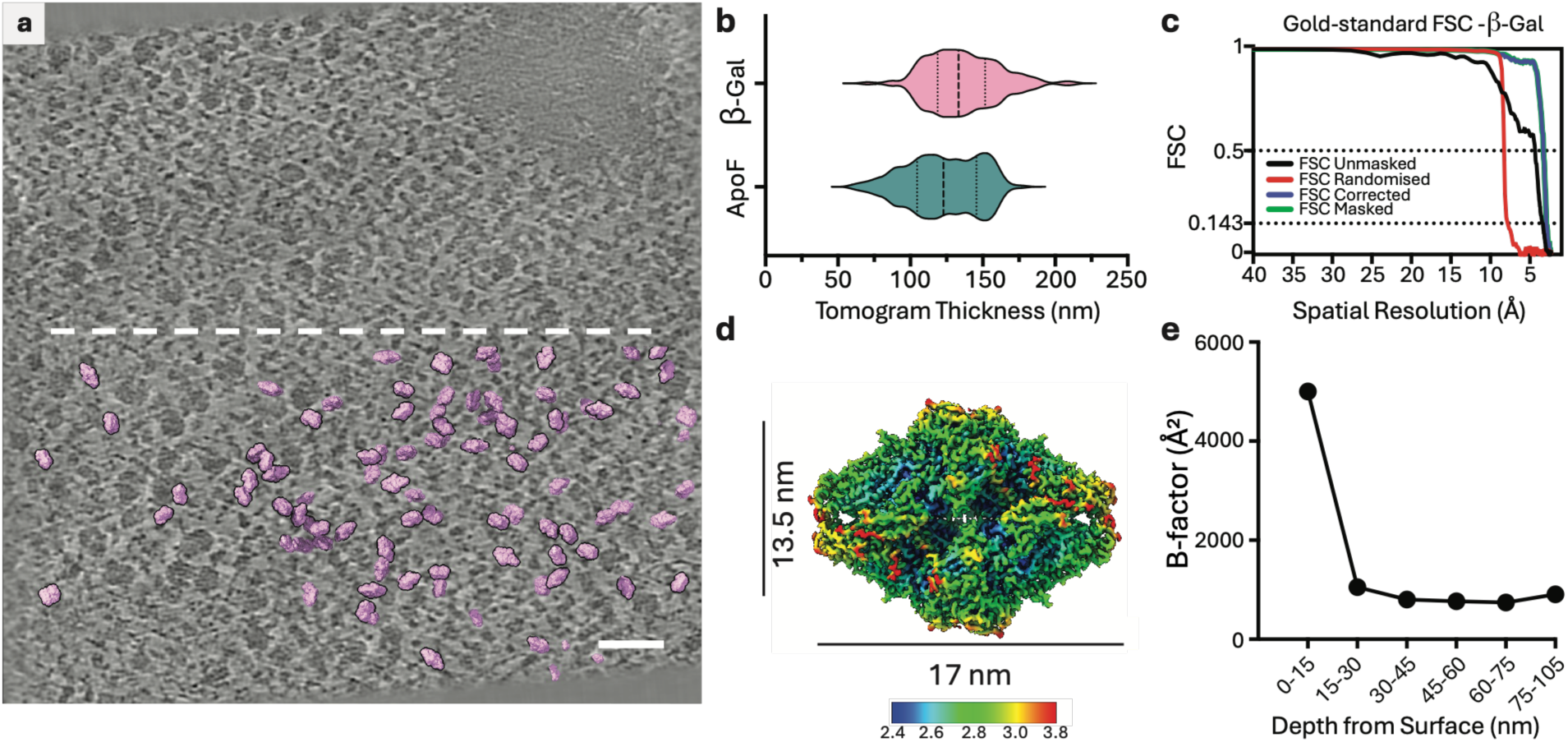
β-Galactosidase as a reporter of cryo-FIB damage in xenon-milled lamellae. a, Representative tomographic slice of a xenon-milled *E. coli* lamella expressing β-galactosidase. Below the dashed line, β-galactosidase volumes (pink) are overlaid at detected particle positions. Scale bar, 50 nm. **b,** Distributions of estimated lamella thickness for xenon-milled *E. coli* expressing β-galactosidase (pink) comparing with the thickness of apoferritin lamellae (cyan). **c,** Gold-standard Fourier shell correlation (FSC) curves for the β-galactosidase reconstruction, showing a global resolution of 2.94 Å at FSC = 0.143. **d,** β-Galactosidase reconstruction colored by local resolution with approximate dimensions of 17 × 13.5 nm. **e,** B-factors estimated by Rosenthal–Henderson analysis of β-galactosidase reconstructions from successive depth intervals. Depth was measured from each particle center to the nearest milled surface. Particles were grouped into 15 nm intervals from 0 to 75 nm, followed by a final 75-105 nm interval.

β-Galactosidase particles were grouped into the same depth intervals used for apoF according to their distance from the nearest milled surface and analyzed. B-factors were highest within 0-15 nm of the surfaces, decreased sharply at 15-30 nm and remained comparatively stable at greater depths (Fig. 5e; Supplementary Fig. 9). Although absolute B-factors were higher than for apoF, β-Galactosidase reproduced the narrow xenon-associated profile despite its more extended, multidomain structure and lower symmetry.

## Discussion

We used recombinant apoF to quantify depth-dependent structural preservation in cellular lamellae. ApoF supported near-atomic resolution structure determination *in situ* and revealed differences in apparent damage depth among four milling workflows. Comparison with endogenous 70S ribosomes in the same xenon-milled tomograms showed that apparent damage depth depends on the reporter and reflects the depth-dependent loss of high-resolution information rather than a direct measurement of ion implantation or a fixed physical boundary.

Purified and *in situ* apoF reached comparable near-atomic resolutions and B-factors despite the cellular lamellae being thicker than the purified specimen. Both reconstructions reached the physical Nyquist limit; rendering EER data at 6K sampling enabled refinement beyond the Nyquist limit (Supplementary Fig. 1). The slope of the Rosenthal-Henderson plot describes how resolution scales with particle number, whereas the intercept depends on per-particle signal and other dataset-specific factors [30], but similar B-factors do not imply equivalent per-particle signal. Purified apoF reached higher resolution for the same number of particles, consistent with a reduced per-particle signal-to-noise ratio in thicker cellular specimens. The comparable slopes nevertheless indicate similar rates of resolution improvement with increasing particle number.

Several factors may have contributed to the near-atomic resolution achieved for apoF *in situ*. The rigidity, structural homogeneity, and octahedral symmetry of apoF facilitate alignment and averaging, while its high intracellular abundance increases particle yield. Similar B-factors may also reflect limitations shared by sample preparation, tomographic acquisition, and processing. Purification included incubation at 70 °C, and exposure to the air-water interface could offset some advantages of thin vitreous ice. Conversely, intracellular environment, including ionic composition and macromolecular crowding may help preserve the apoF cage. These possibilities remain untested, but our results demonstrate that near-atomic-resolution structure determination is achievable *in situ* for an abundant, structurally tractable target.

Within the xenon dataset, resolution improved and B-factors decreased from the 60-100 nm group to the 125-150 nm group, with both trends reversing in the 150-175 nm group (Supplementary Fig. 3). Lower resolution in the thinnest group may reflect a higher proportion of particles affected by milling-induced damage, as surface-proximal regions occupy a greater fraction of the specimen volume in thinner lamellae. In the thickest group, thickness dependent reduction in per-particle signal may contribute to the lower resolution. The observed trend is consistent with competing effects of surface-associated damage and specimen thickness, although their relative contributions were not quantified. A previous large-scale *in situ* ribosome analysis also found increased B-factors in lamellae thicker than 225 nm [10]. The thickness at which resolution declines may depend on the specimen, molecular target, and experimental workflow.

Using apoF, we estimated apparent damage depths of approximately 15 nm for xenon, 30 nm for gallium, 45 nm for argon, and at least 75 nm for oxygen (Fig. 3). Xenon produced the thinnest lamellae and oxygen the thickest, with gallium and argon producing intermediate thicknesses (Fig. 2). Mean thickness does not explain all differences: gallium and argon-milled lamellae had similar mean thicknesses but different B-factors. Excluding particles within 75 nm of oxygen-milled surfaces improved reconstruction quality despite reducing particle number (Supplementary Fig. 4). The ordering of apparent damage depths is qualitatively consistent with species-dependent ion penetration but does not establish a simple relationship with ion mass, as it also depends on energy transfer through atomic collisions, electronic excitation and ionization, incidence angle, ion fluence, and chemical interactions. Therefore, B-factors reflect high-resolution signal attenuation rather than directly measuring ion implantation or physical damage depth.

Species-dependent milling rates may also contribute to differences in final lamella thickness. In vitrified biological specimens, milling rates depend on both ion species and incidence angle and cannot be predicted from ion mass alone. Under previously examined conditions, xenon milled most rapidly, whereas argon milled more slowly than oxygen [31]. Sputter yield and surface texturing also vary with target material [31], while plasma gases produce different probe profiles and curtaining behavior. Oxygen plasma can contain O⁺ and O₂⁺ [32]; and mass-to-charge characterization of a Helios Hydra PFIB detected Ar²⁺ at an argon-source operating power of 200 W [32, 33]. We did not characterize beam composition, and equivalent aperture settings do not ensure equivalent currents, current-density distributions or ion fluences. Our results therefore compare the performance of complete milling workflows rather than isolate intrinsic effects of ion species.

Disentangling the effects of ion species, beam energy and polishing conditions would require controlled comparisons accounting for current density, ion fluence, incidence angle, and final lamella thickness, although matching these parameters simultaneously may be difficult with current instrumentation. Low-voltage polishing can reduce apparent damage in gallium-milled lamellae, but larger ion probes at lower voltages can complicate the preparation of thin lamellae and increase curtaining [9, 15]. These limitations are relevant to plasma FIB workflows, where maintaining well-focused xenon probes at lower accelerating voltages can be challenging [17]. We did not test low-voltage polishing in this study, and whether it further reduces the apparent damage observed with each ion species remains to be determined.

Within the same lamellae, apoF indicated an apparent xenon-associated damage depth of approximately 15 nm, whereas ribosome B-factors remained elevated throughout the first 30 nm and decreased only in the 30-45 nm interval (Fig. 4), consistent with previous measurements [8–10, 14, 16]. The 25-30 nm ribosome dimensions limit the spatial precision of depth assignment from its center. Differences in composition, architecture, conformational heterogeneity, and susceptibility to structural perturbation may also contribute. Damage may affect template matching, alignment and reconstruction differently for each target. Some targets may also benefit more from symmetry averaging, producing different apparent profiles even when the underlying physical damage is similar.

β-galactosidase has a comparable molecular mass to apoF but a more extended, multidomain structure and lower symmetry. It nevertheless showed the strongest loss of high-resolution information within approximately 15 nm of xenon-milled surfaces, consistent with the apoF profile (Fig. 5). Differences in absolute B-factors among all three reporters emphasize that these values remain reporter-dependent. The β-galactosidase reconstruction also shows that recombinant bacterial expression and xenon milling support high-resolution *in situ* structure determination beyond highly symmetric apoF.

Applying ribosome-based exclusion distances of approximately 30-45 nm from each milled surface to other targets could discard particles useful for STA. For example, in a 128 nm thick lamella (the xenon mean), excluding 15 nm from each surface leaves approximately 98 nm of lamella thickness, compared with 68 nm or 38 nm for 30 nm or 45 nm exclusions. These values describe the thickness retained after exclusion and do not establish that the retained region is undamaged. The appropriate exclusion distance depends on the target, milling workflow, and desired resolution. Nevertheless, avoiding unnecessarily conservative surface exclusion will increase particle yield, particularly for low-abundance complexes.

Recombinantly expressed, structurally tractable proteins provide a framework for benchmarking cryo-FIB preparation and pursuing high-resolution structure determination *in situ*. Extending this approach to reporters with diverse sizes, compositions, symmetries and architectures could help resolve how beam parameters, polishing conditions and specimen type influence structural preservation. The apoF and β-galactosidase datasets demonstrate that increased intracellular abundance can supply sufficient particles for high-resolution subtomogram averaging in *E. coli*, including for a lower-symmetry, multidomain complex. Controlled expression in mammalian cells could extend this strategy to proteins requiring a eukaryotic environment, provided that native-like assembly, stoichiometry, localization and function are verified. Together, our findings support evaluating cryo-FIB workflows according to the molecular target and desired resolution, with usable lamella volume determined by the combined constraints of target-dependent preservation, specimen thickness and particle availability.

## Materials and Methods

### Recombinant apoferritin expression and purification

The hFerritinH expression plasmid was a gift from Trevor Douglas (Addgene plasmid no.122652; http://n2t.net/addgene:122652; RRID: Addgene 122652). The plasmid was transformed into *Escherichia coli* BL21 Star™ (DE3) cells. An overnight culture was diluted 1:100 into 500 mL of 2×YT medium supplemented with kanamycin (50 μg/mL), and the culture was grown at 37 °C with shaking at 220 r.p.m. When the culture reached an OD₆₀₀ of 0.5, protein expression was induced with 0.5 mM IPTG, and incubation was continued for 3 h at 37 °C. Cells from a 2-mL aliquot of the induced culture were collected by centrifugation, washed with phosphate-buffered saline (PBS), and resuspended in either 50 or 100 µL PBS for immediate preparation of EM grids. The remainder of the culture was harvested by centrifugation, and the pellet was stored at −80 °C until purification.

Cell pellets were resuspended in 25 mL of buffer A (30 mM HEPES, pH 7.5, 300 mM NaCl, 1 mM MgSO₄) supplemented with 1 mg/mL lysozyme and one protease inhibitor cocktail tablet (Roche). The suspension was incubated for 15 min at 4 °C with rotation and then lysed by sonication. Cell debris was removed by centrifugation for at 20,000 × g for 30 min at 4 °C. The clarified supernatant was heated to 70 °C for 10 min while stirring at 100 rpm, cooled on ice, and the resulting milky solution was centrifuged at 20,000 × g for 30 min at 4 °C. Ammonium sulfate was added to the supernatant to approximately 52.5% saturation, and sample was stirred for 10 min at 4 °C. The mixture was centrifuged at 14,000 × g for 10 min at 4 °C. The resulting pellet was resuspended in 6 mL of buffer B (20 mM HEPES, pH 7.5, 300 mM NaCl) and dialyzed overnight against 2 L of buffer B using a 10 kDa MWCO membrane. Further purification was performed by size-exclusion chromatography on a Superose 6 10/300 GL column (GE Healthcare) equilibrated with 20 mM HEPES, pH 7.5, 300 mM NaCl, and 1 mM TCEP. Fractions corresponding to the 24-mer assembly were pooled and concentrated to 4-8 mg/mL using a centrifugal concentrator with a 100 kDa MWCO. Purified protein was used immediately to prepare cryo-EM grids.

### Recombinant β-galactosidase expression

Plasmid pSG 25 was a gift from Albert Dahlberg (Addgene plasmid no. 63867; RRID: Addgene 63867) and was received as a bacterial stab. Bacteria from the stab were streaked onto tetracycline selective agar, and a single colony was picked and cultured overnight at 37 °C with shaking at 220 r.p.m. in 5 mL of LB medium containing tetracycline (10 μg/mL). Plasmid DNA was isolated from the overnight culture and subsequently transformed into *E. coli* BL21(DE3) cells (Thermo Fisher Scientific, catalog no. ECO114). Transformed cells were grown overnight at 37°C with shaking at 220 r.p.m. in 5 mL LB medium containing tetracycline (10 μg/mL). A 0.5 mL aliquot of the overnight culture was used to inoculate 25 mL of LB medium containing tetracycline (10 μg/mL) and grown at 37°C with shaking at 220 r.p.m. until the culture reached an OD600 of approximately 0.6. Protein expression was induced in a 5 mL aliquot by adding IPTG to a final concentration of 0.4 mM, and the culture was incubated for 3 h at 37°C with shaking at 220 r.p.m. Cells from 2 mL of the induced culture were collected by centrifugation at 2000 x g for 3 min, and the pellet was resuspended in 75 μL of PBS for cryo-EM grid preparation.

### Cryo-EM grid preparation

Purified apoF (3.5 μl) was applied to Quantifoil R1.2/1.3 Cu 300-mesh holey carbon grids (Quantifoil Micro Tools GmbH, Germany). Grids were blotted for 3s at 10°C and 95% relative humidity and then plunge frozen in liquid ethane using a Vitrobot mark IV (Thermo Fisher Scientific).

For *in situ* apoF and β-galactosidase samples, the corresponding cell suspensions (4 μl) were applied to Quantifoil R1.2/1.3 Cu 300-mesh holey carbon grids (Quantifoil Micro Tools GmbH, Germany). Grids were back-blotted for 6s at 22°C and 95% relative humidity and then plunge frozen in liquid ethane using a Leica EM GP2 (Leica Microsystems).

### Cryo-FIB milling workflows

#### Xenon milling

Vitrified grids were clipped into TomoGrid (Thermo Fisher Scientific), and cryo-lamellae were prepared using an Arctis Plasma FIB-SEM (Thermo Fisher Scientific) using MAPS (v3.36) and AutoTEM Cryo software (v2.4.6) (Thermo Fisher Scientific). Prior to milling, the grids were sputter-coated with platinum for 2 min at 70 nA, followed by organometallic platinum deposition for 45 s using the gas injection system (GIS) and a second platinum sputter coating for 2 minutes at 70 nA.

Automated lamella preparation was performed with a xenon plasma ion beam operated at an accelerating voltage of 30 kV. Stress-relief cuts were first milled on either side of the intended lamella, and a milling angle of 12° was used for all lamellae. Lamellae were milled to a nominal width of approximately 10-12 µm using three sequential milling steps to reach an intermediate thickness of approximately 300 nm, followed by polishing. Rough milling was performed using rectangle patterns at 1 nA with a pattern offset of 1 µm. Medium milling was performed using regular cross-section patterns at 0.3 nA beam current with a pattern offset of 600 nm. Fine milling was performed using regular cross-section patterns at 0.1 nA with a pattern offset of 300 nm. Final thinning was performed at 30 pA, using cleaning cross-section patterns with a target thickness of 120 nm.

#### Xenon coarse milling followed by argon or oxygen final thinning

Vitrified grids were clipped into AutoGrid (Thermo Fisher Scientific), and cryo-lamellae were prepared using an Arctis Plasma FIB-SEM (Thermo Fisher Scientific) operated with MAPS (v3.36) and AutoTEM Cryo software (v2.4.6) (Thermo Fisher Scientific). Prior to milling, the grids were sputter-coated with platinum for 2 min at 70 nA, followed by organometallic platinum deposition for 80 s using the GIS and a second platinum sputter coating for 2 min at 70 nA.

Automated lamella preparation included automated eucentric height correction, beam coincidence alignment, reference definition, autofocus, and milling-angle optimization prior to patterning. Lamellae were prepared with nominal dimensions of 12 μm × 3 μm and a target final thickness of 70 nm. Stress-relief cuts were first milled on either side of the intended lamella, and a milling angle of 12° was used for all lamellae.

For both workflows, a xenon plasma ion beam operated at an accelerating voltage of 30 kV was used for rough and medium milling. After the medium milling step, the plasma source was switched to either argon or oxygen for fine milling and polishing steps. Using a common xenon-based coarse milling protocol ensured identical bulk material removal prior to the final thinning stage.

Rough milling was performed using rectangle patterns at 1 nA, with a 1 μm pattern offset. Medium milling was performed using regular cross-section patterns at 0.3 nA with a 600 nm pattern offset. Fine milling was performed using regular cross-section patterns at 60 pA (argon) and 90 pA (oxygen) with a 300 nm pattern offset. Final thinning was performed using cleaning cross-section patterns. The first polishing step was performed at 20 pA with a 50 nm pattern offset, followed by two or three iterations of a second polishing step at 20 pA with zero pattern offset to a target thickness of approximately 70 nm. Throughout automated milling, reference-based drift correction, autofocus, and low-dose SEM imaging were used to maintain accurate pattern placement and monitor lamella quality.

### Gallium milling

Vitrified grids were clipped and cryo-lamellae were prepared using an Aquilos2 FIB-SEM (Thermo Fisher Scientific) using MAPS (v3.36) and AutoTEM Cryo software (v2.4.6) (Thermo Fisher Scientific). Prior to milling, the grids were sputter-coated with platinum for 15 seconds at 30 mA, followed by organometallic platinum deposition for 35 s using the gas injection system (GIS) and a second platinum sputter coating for 15 seconds at 30 mA.

Automated lamella preparation was performed with a gallium ion beam operated at an accelerating voltage of 30 kV. Stress-relief cuts were first milled on either side of the intended lamella, and a milling angle of 12° was used for all lamellae. Lamellae were milled to a nominal width of approximately 10-12 µm using three sequential milling steps to reach an intermediate thickness of approximately 300 nm, followed by polishing. Rough milling was performed using rectangle patterns at 1 nA with a pattern offset of 1 µm. Medium milling was performed using regular cross-section patterns at 0.5 nA beam current with a pattern offset of 600 nm. Fine milling was performed using regular cross-section patterns at 0.3 nA with a pattern offset of 300 nm. Final thinning was performed at 30 pA, using cleaning cross-section patterns with a target thickness of 90 nm.

### Cryo-electron tomography data acquisition

Tilt series were acquired on a Titan Krios G4 or Krios 5 transmission electron microscopes (Thermo Fisher Scientific) operating at an accelerating voltage of 300 kV and equipped with a Selectris X energy filter and a Falcon 4i direct electron detector. Data collection was performed using Tomography 5 software (Thermo Fisher Scientific) with a dose-symmetric tilt scheme and a 10 eV energy slit. Tilt series were acquired over an angular range of −54° to +54° starting from the pre-determined milling angle with a tilt increment of 3°.

Movies were recorded in counting mode and saved in electron event representation (EER) format at a nominal magnification of 105,000x, corresponding to a calibrated pixel size of 1.19 Å/pixel. The target defocus range was −3 µm to −1 µm. Each tilt image was recorded as an EER movie, resulting in a target dose range of 3.5-4 e⁻/Å² per tilt and a total accumulated dose of approximately 130-150 e⁻/Å² per tilt-series. Automated acquisition, including stage tilting, tracking, focusing, and dose management, was controlled by Tomography 5 software (Thermo Fisher Scientific).

Data were collected either using aberration-free image shift (Multishot) with a maximum beam-image shift of 3 µm or without image shift (single-position). At comparable lamella thicknesses, apoF reconstruction resolutions were similar between the two acquisition modes (Supplementary Fig. 10).

### Processing of purified apoF data

Movie frames were preprocessed using Warp v2.0.0dev33 [34], including gain correction, motion correction, dose weighting, per-tilt contrast transfer function (CTF) estimation using a tilt-aware CTF model, and generation of tilt stacks. Motion correction was done using 8K EER movies, and Fourier cropped to 4K with a pixel size of 1.19 Å. The resulting tilt series were aligned using patch tracking in AreTomo2 [35] at a binning factor of 8 and reconstruction by weighted back-projection (WBP) in Warp. ApoF particle coordinates and best-scoring template-matching orientations were determined using pytom-match-pick [25], with a low-pass filtered, resampled version of EMD-15854 as the reference. Subtomograms were extracted in Warp at a binning factor of 4, with a 64³ voxel box size. An initial reference volume was generated using the relion_reconstruct program in RELION 4.0 and used for 3D auto-refinement with octahedral (O) symmetry imposed, resulting in a 9.83 Å reconstruction.

The refined particle set was then transferred to M [36] for high-resolution refinement using the unbinned 4K-sampled data (1.19 Å/pixel). Seven iterative refinement rounds were performed to optimize particle poses and motion trajectories together with per-tilt CTF and per-series optical parameters, including beam tilt and higher-order aberrations. This refinement produced a 2.4 Å apoF reconstruction, approaching the physical Nyquist limit. To enable refinement beyond the physical Nyquist limit imposed by 1.19 A physical pixel sampling, the EER data were resampled in Warp at 6K sampling, corresponding to 0.79 Å per pixel, and the particles were re-extracted. Further high-resolution refinement in M yielded a final 1.97 Å apoF reconstruction that extended beyond the physical Nyquist limit of 4K sampling

Global resolution was estimated from the gold standard Fourier shell correlation (FSC) between independently refined half-maps using the 0.143 criterion. Final density maps were inspected and visualized using UCSF Chimera or ChimeraX [37].

### Processing *in situ* apoF data

Gain correction, motion correction CTF estimation and tilt-stack generation were performed in Warp, with raw EER data grouped in sets of six frames. Tilt series alignment was carried out using AreTomo2 at a binning factor of 8 and were subsequently reconstructed in Warp. ApoF particles were initially picked using pytom-match-pick, with a low-pass filtered, resampled version of EMD-15854 as the reference, yielding 150,580 picked particles. Subtomograms were exported in Warp at bin 4 with a 64³ voxel box size, and reference volumes were generated using RELION 4.0. RELION 3D auto-refinement with O symmetry imposed, produces a 9.83 Å map. Refined particles were resampled using MTools to bin 1 (1.19 Å/pixel). Subsequent pose refinement was performed in M over seven iterative rounds, leading to a 2.4 Å reconstruction, approaching the 2.38 Å physical Nyquist limit.

A second data set was collected to increase particle numbers for depth-dependent analysis of structure preservation after milling. This dataset comprised 150 additional tomograms and yielded additional 115,707 particles. These particles were combined with the initial dataset, yielding a total of 266,287 particles. Refinement in RELION followed by pose refinement in M using the 6K-sampled data yielded a final 2.05 Å apoF reconstruction.

### Processing of *in situ* ribosome data

Endogenous *E. coli* 70S ribosomes were identified by template matching using pytom-match-pick, with a low-pass filtered, resampled version of EMD-28254 [38] as the reference. This yielded approximately 92,160 candidate particles, that were imported into Warp for subtomogram extraction at bin 2 using a 224³ voxel box size. A round of 3D-auto refinement followed by 3D classification was run in RELION using a soft mask of the 50S subunit. Four classes were selected and merged, yielding 70,087 particles for a final run of refinement, and M post-processing, achieving a 3.89 Å reconstruction.

To further improve this map and resolve ribosome heterogeneity, the particles were analyzed using tomoDRGN [39]. M-aligned particles were extracted in Warp at bin 4, and analyzed through tomoDRGN’s variational auto-encoder, to further segment heterogeneous 70S particles (zdim 128). This yielded a UMAP with 3 distinct groups, one containing 50S alone, one with 70S alone, and a third group with broken or junk particles. The 70S and 50S subsets contained 16,627 and 14,449 particles, respectively. Each subset underwent a final round of RELION 3D auto-refinement followed by multi-species refinement in M, with apoF included as an additional species. This yielded final 70S and 50S reconstructions at 3.26 Å and 3.6 Å resolution, respectively.

### Processing *in situ* β-galactosidase data

Gain correction, motion correction CTF estimation and tilt-stack generation were performed in Warp v2.0.0dev33, with raw EER data grouped in sets of nine frames. Tilt series alignment was carried out at binning factor 8 using AreTomo2, and the aligned data were subsequently reconstructed back in Warp. β-galactosidase particles were initially picked using pytom-match-pick, with a low-pass filtered, resized version of EMD-5995 [26] as the reference. 3D particles were exported in Warp at bin 4 with a 112³ voxel box size, and reference volumes were generated using RELION 5.0 [40]. Next, RELION 3D auto-refinement was run, applying D2 symmetry, to produce a 9.69 Å map. 3D classification was performed in RELION v5.0.1. An initial set of 34,191 particles was processed with six output groups using an initial reference low-pass filtered to 25 Å and with C1 symmetry. The particle mask diameter was set to 220 Å. CTF correction was enabled, and the run was performed for 50 iterations with the regularisation fudge parameter set to 0.5. Angular sampling used HEALPix order 2 with oversampling set to 1, and translational searches used an origin-offset range of 5 pixels with 2-pixel sampling. The standard deviation for local angular searches over the Euler angles was set to 7.5°. Solvent flattening and zero masking were applied using a solvent mask. Following classification, selected output groups were pooled, yielding 24,893 particles for downstream refinement. Refined particles were resampled using MTools to bin 1 (1.19 Å/pixel). Subsequent pose refinement was performed in M over five iterative rounds, leading to a 2.94 Å reconstruction.

### Binning particles according to distance from the nearest lamella surface

To assign apoF and ribosome particles to depth bins, we wrote a custom Python script that reads particle-center coordinates from the pytom-match-pick STAR files and calculated their distance from the upper and lower milled surfaces. For each particle, the distance along the tomogram z axis between its z coordinate and each interpolated surface at the corresponding x–y position was calculated, and the smaller of the two values was retained. Boundary models were manually annotated using IMOD and converted into text files for input to the Python script. Overall lamella thickness was estimated using IMOD boundary models loaded into Slabify (https://github.com/CellArchLab/slabify-et). Particles centers were then assigned to consecutive 15 nm distance bins beginning at either milled surface (0-15 nm, 15-30 nm and so forth) and extending to the lamella midplane with a final bin of 75-105 nm. Particles from the upper and lower halves of each lamella were pooled according to their distance from the nearer surface. Bin assignments were visually validated by mapping the particles back into the tomograms in ArtiaX [41].

For B-factor estimation, the STAR file corresponding to each depth bin further split into groups of increasing particle number for B-factor estimation. Particles were exported in Warp at bin 4, and reference volumes were generated using relion_reconstruct for each particle bin, followed by 3D auto-refinement. Particles were then imported back into Warp-M at bin 1 and run through MCore without any refinements. B-factors calculated based on the linear fit of resolution-2 vs ln (number of particles).

### LCCmax comparison between ion species data sets

To quantify the differences in LCCmax values between particles picked from lamellae milled with gallium, oxygen, argon, or xenon, we used a custom python script to separate particles from STAR files generated after 3DTM with pytom-match-pick into distinct bins based on their location within the tomograms. We used a modified version of the python script used for sorting particles into 15 nm bins for the B-factor analysis. For the LCC comparison, we elected to bin particles within the 0-45 nm and 45-105 nm into separate bins. Particles were then organized based on their LCCmax values, and the top 1500 scoring particles from each data set were extracted and analyzed in GraphPad prism.

## Acknowledgments

We thank Ricardo Diogo Righetto and Davide Tamborrini (University of Basel) for help with PyTOM, WarpTools and Slabify, and Martin Obr (Thermo Fisher Scientific) for advice on RELION processing. We thank Bryan Sibert (St. Jude Children’s Research Hospital) for processing advice and assistance with 3D classification of 70S ribosomes using TomoDRGN, and Walid Abu Al-Afia (St. Jude Children’s Research Hospital) for computational support. We also thank Jarrad Weirich (St. Jude Children’s Research Hospital) for configuring remote access to the Arctis and Krios G4 microscopes, and Oliver Raschdorf and Ron Kelley (Thermo Fisher Scientific) for microscope access and discussions. We are grateful to Radostin Danev (The University of Tokyo), Chris Russo (MRC Laboratory of Molecular Biology, Cambridge) and Peijun Zhang (Electron Bio-Imaging Centre, Diamond Light Source, and University of Oxford) for helpful discussions. We also thank the Cryo-EM Center and its staff in the Department of Structural Biology at St. Jude Children’s Research Hospital and Center of Excellence for Structural Cell Biology staff for providing resources and support.

## Author Contributions

A.K. conceived the idea and supervised the project with GS. R.M.N., M.Z.Q. and A.K. designed the experiments, performed FIB milling and acquired TEM data. R.M.N. and M.Z.Q. processed the xenon, gallium, argon, and oxygen datasets. D.K. and AK performed FIB milling and acquired TEM data for the xenon dataset at Thermo Fisher Scientific, Eindhoven. M.Z.Q. vitrified samples and clipped grids. V.M. cultured *E. coli* and expressed β-galactosidase. V.M. and J.M.d.l.R.T. processed the β-galactosidase dataset. A.R. cultured *E. coli* and expressed and purified apoF. Y.S. performed initial milling on the Zeiss Crossbeam and supported TEM and Aquilos workflows. H.Y. performed initial experiments using the Aquilos 2. C.T. supported plasma- and gallium-FIB milling. S.A.H. provided operational support and oversight for TEMs. S.S. provided operational support for FIB instruments and contributed to discussions of the figures. J.M.d.l.R.T. provided computational support for software and hardware. R.K. provided operational support and oversight for cell culture and protein purification. R.M.N., M.Z.Q., G.S. and A.K. analyzed the data and prepared the manuscript with contributions from all authors.

## Competing interests

Dimple Karia is an employee of Thermo Fisher Scientific. The other authors declare no competing interests.

## Data availability

The subtomogram averaged maps have been deposited in the Electron Microscopy Data Bank (EMDB) under accession codes [EMD-XXXXX] (apoF, xenon), [EMD-XXXXX] (apoF, argon), [EMD-XXXXX] (apoF, gallium), [EMD-XXXXX] (apoF, oxygen), [EMD-XXXXX] (70S ribosome, xenon) and [EMD-XXXXX] (β-galactosidase, xenon). Ion species indicate the milling conditions used to prepare the cellular lamellae. The oxygen-milled apoF map was reconstructed from the subset of particles remaining after exclusion of particles within 75 nm of either milled surface.

Raw electron microscopy data will be available at the Electron Microscopy Public Image Archive (EMPIAR).

---

**Supplementary Table 1.** Data collection parameters, and processing outcomes for Argon, Gallium, Oxygen, Xenon, and Purified data sets. Particle numbers, resolutions, and B-factors are generated from apoferritin.

**Figure 5**
|  | Argon | Gallium | Oxygen | Xenon | Purified |
| --- | --- | --- | --- | --- | --- |
| Microscope | Krios G4 | Krios G4 | Krios G4 | Krios G4 | Krios G4 |
| Voltage (kV) | 300 | 300 | 300 | 300 | 300 |
| Detector | Falcon 4i | Falcon 4i | Falcon 4i | Falcon 4i | Falcon 4i |
| Energy Filter | Selectris X | Selectris X | Selectris X | Selectris X | Selectris X |
| Magnification | 105000x | 105000x | 105000x | 105000x | 105000x |
| Pixel Size (Å) | 1.19 | 1.19 | 1.19 | 1.19 | 1.19 |
| Defocus Range (µm) | -3 to -1 | -3 to -1 | -3 to -1 | -3 to -1 | -3 to -1 |
| Total Tomograms | 494 | 102 | 256 | 447 | 55 |
| Tomograms Used | 238 | 82 | 197 | 289 | 55 |
| Number of Particles | 83160 | 79256 | 105087 | 266,287 | 67,739 |
| Resolution (Å) | 2.88 | 2.76 | 6.10* (3.6 from 45,438 particles from 75-105 nm layer) | 2.05 | 1.97 |
| B-factors (Å<sup>2</sup>) | 153.9 | 128.2 | 1250 | 122.3 | 123.5 |

**Supplementary Figure 1.**
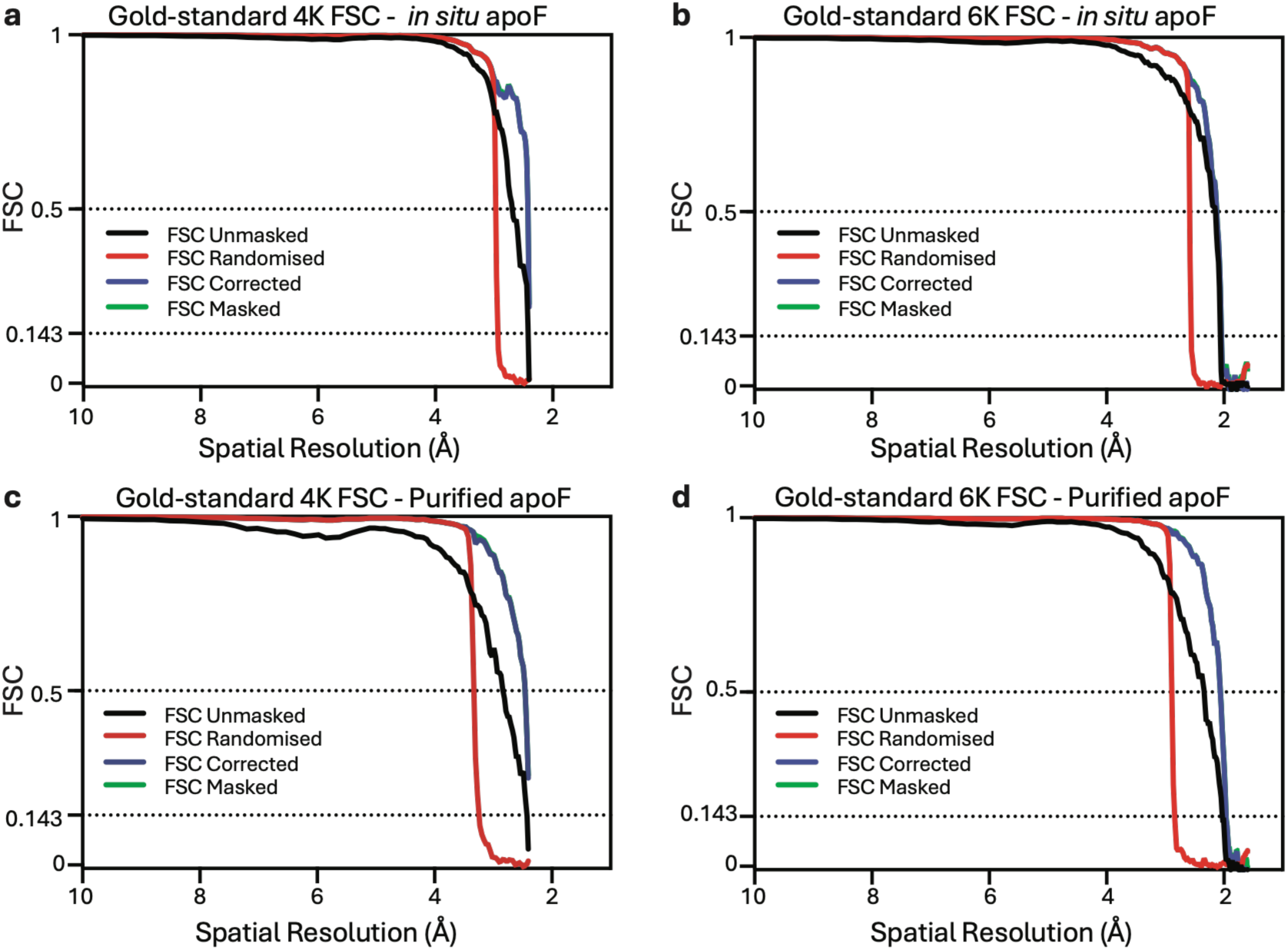
Apoferritin refinement beyond the physical Nyquist limit. a, Gold-standard Fourier shell correlation (FSC) curves for *in situ* apoferritin from xenon-milled lamellae, reconstructed from EER data rendered at 4K sampling and reaching the physical Nyquist limit of 2.38 Å. **b,** Corresponding FSC curves after processing at 6K sampling, yielding a resolution of 2.05 Å, beyond the physical Nyquist limit. **c,** Gold-standard FSC curves for purified apoferritin reconstructed from EER data rendered at 4K sampling, reaching the physical Nyquist limit of 2.38 Å. **d,** Corresponding FSC curves after processing at 6K sampling, yielding a resolution of 1.97 Å, beyond the physical Nyquist limit. Resolutions in **b** and **d** were determined at FSC = 0.143.

**Supplementary Figure 2.**
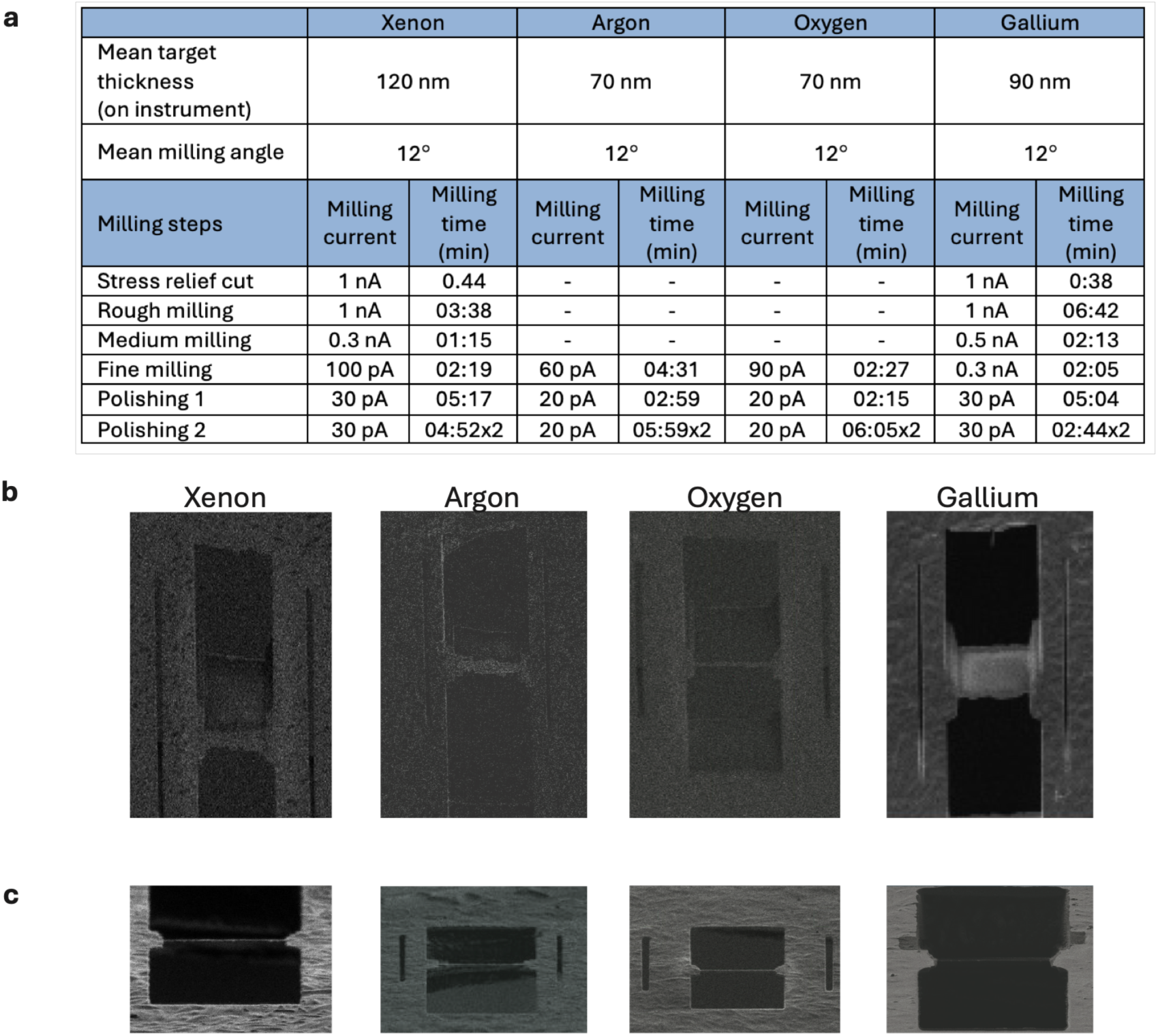
FIB-milling parameters for preparation of *E. coli* lamellae. a, Instrument target thicknesses, milling angles, ion currents and approximate times for each milling step in the xenon, argon, oxygen and gallium workflows using AutoTEM Cryo. Plasma FIB lamellae were initially milled with xenon to approximately 600 nm before final thinning and polishing with xenon, argon or oxygen. Representative SEM (b) and FIB (c) images of lamella sites prepared using xenon, argon, oxygen and gallium.

**Supplementary Figure 3.**
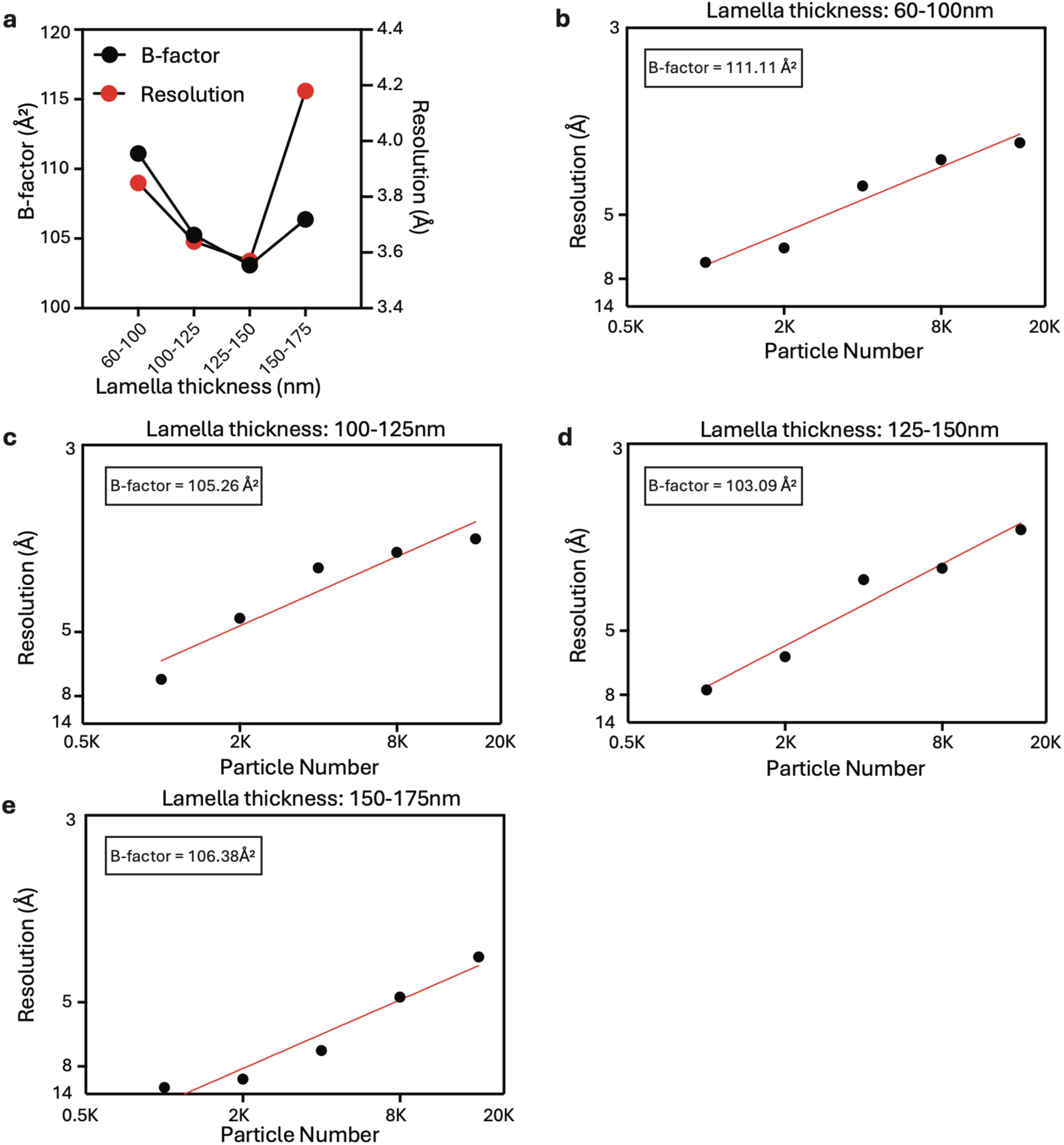
Effect of lamella thickness on apoferritin B-factors and resolution. a, B-factors (black, left axis) and reconstruction resolutions (red, right axis) for *in situ* apoferritin from xenon-milled *E. coli* lamellae grouped by thickness. **b-e,** Rosenthal– Henderson analyses for particles from lamellae 60-100 nm (**b**), 100-125 nm (**c**), 125-150 nm (**d**) and 150-175 nm (**e**) thick.

**Supplementary Figure 4.**
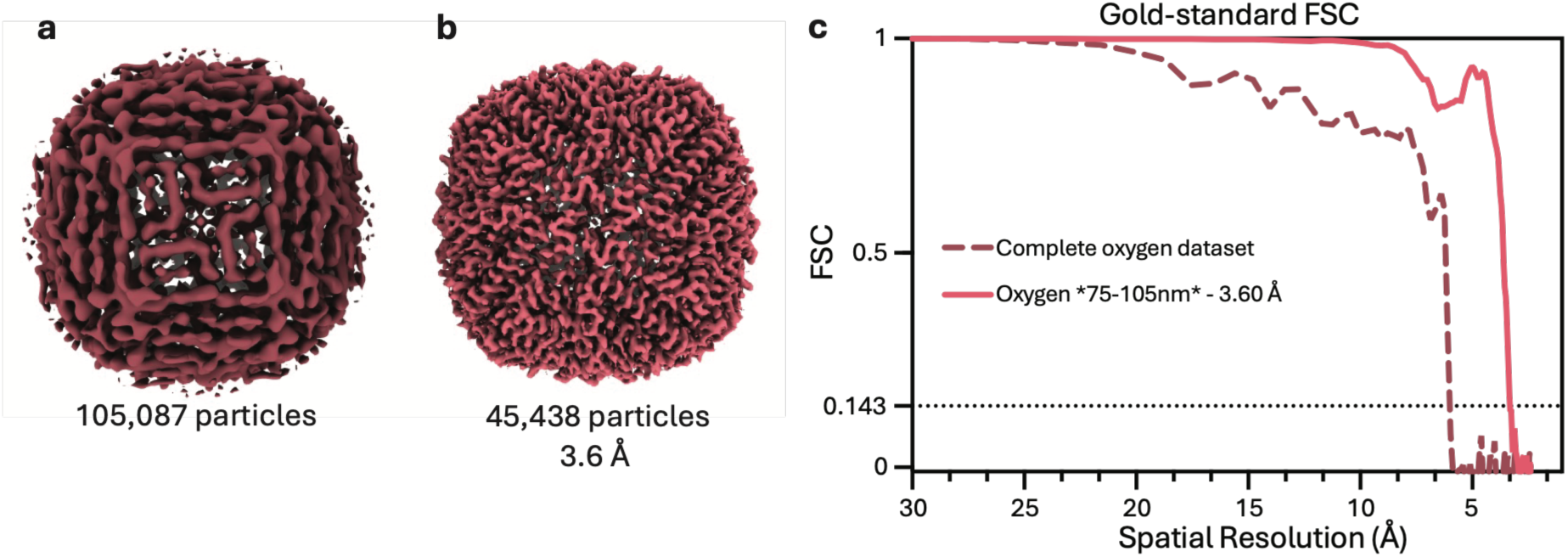
Excluding surface-proximal particles improves apoferritin reconstruction from oxygen-milled lamellae. a, Apoferritin reconstruction after M refinement using the complete oxygen PFIB dataset (105,087 particles). b, Corresponding reconstruction using 45,438 particles whose centers were located 75–105 nm from the nearest milled surface, following exclusion of particles within 75 nm. Both maps are displayed at 3σ. c, Gold-standard Fourier shell correlation (FSC) curves for the complete dataset (dashed line) and retained particle subset (solid line). The retained subset yielded a resolution of 3.60 Å at FSC = 0.143. The complete-dataset reconstruction did not reproduce the expected apoferritin subunit arrangement, consistent with unreliable particle alignment.

**Supplementary Figure 5.**
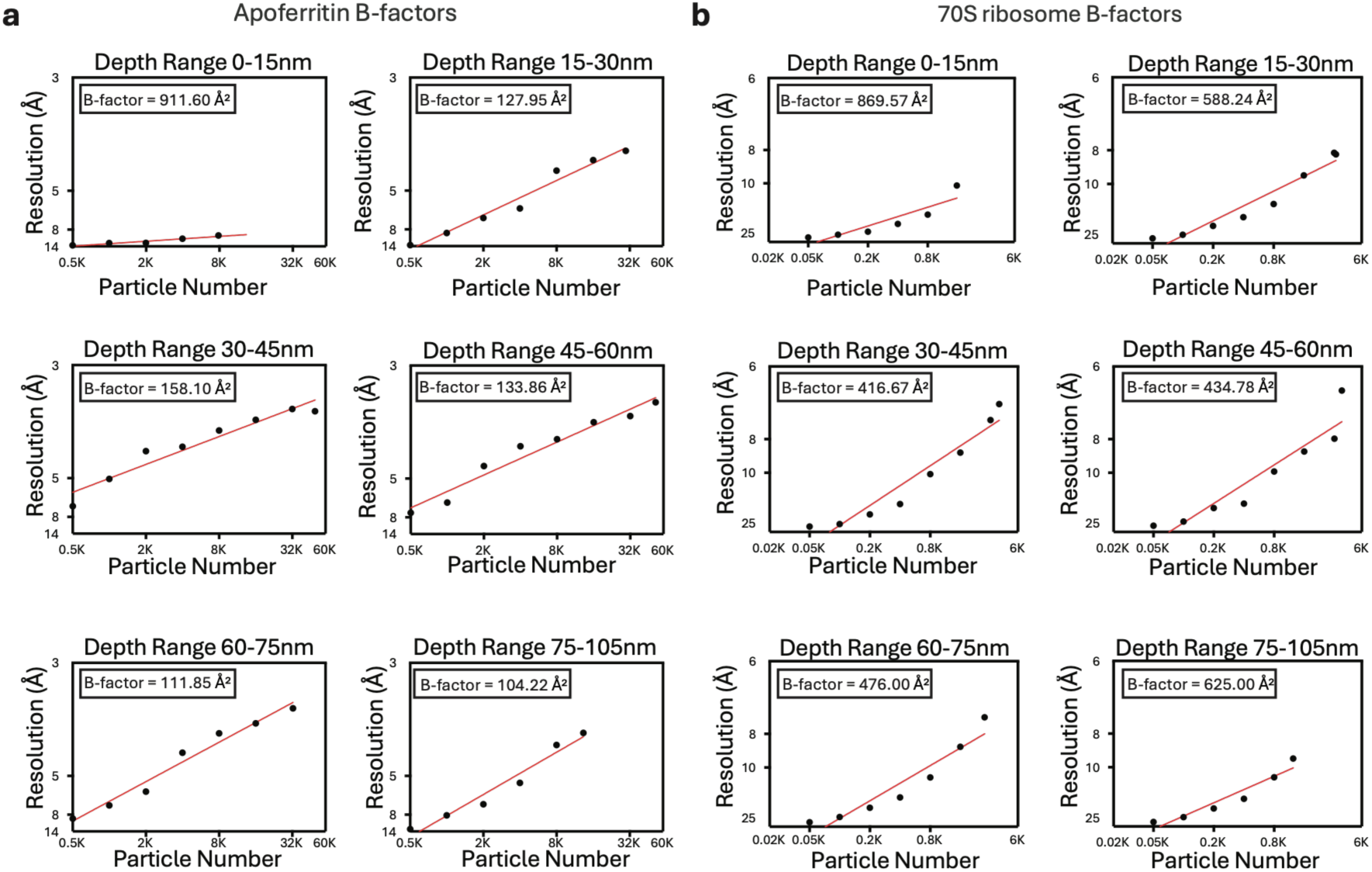
Rosenthal-Henderson analyses of apoferritin and 70S ribosomes in xenon-milled lamellae. a, b, Rosenthal-Henderson plots for *in situ* apoferritin (**a**) and 70S ribosomes (**b**) from the same xenon-milled *E. coli* tomograms, grouped by particle depth. Depth was measured from each particle center to the nearest milled surface. Particles were grouped into 15 nm intervals up to 75 nm, followed by a final 75-105 nm interval.

**Supplementary Figure 6.**
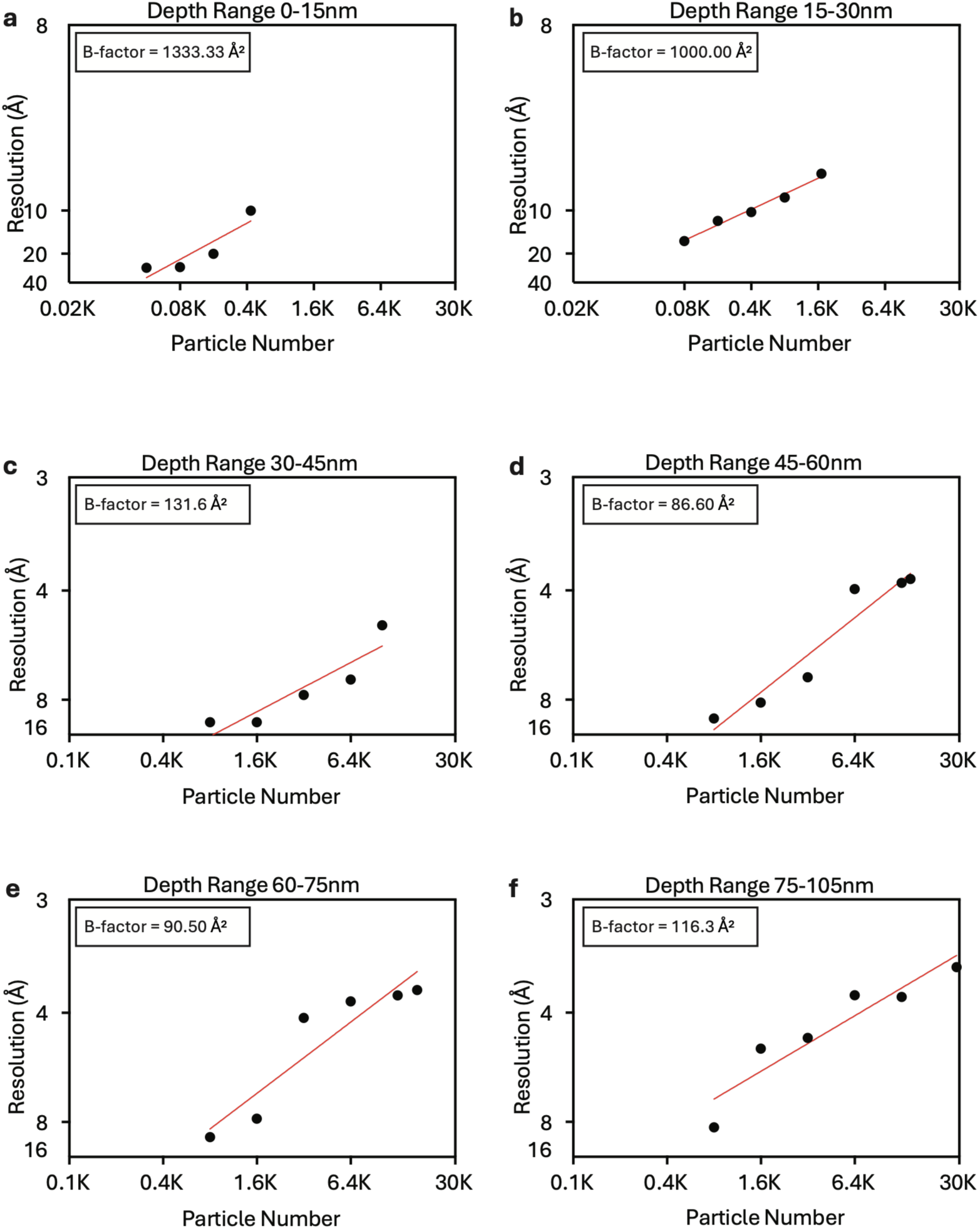
Rosenthal-Henderson analyses of apoferritin in gallium-milled lamellae. a-f, Rosenthal-Henderson plots for *in situ* apoferritin from gallium-milled *E. coli* lamellae, grouped by particle depth: 0-15 nm (**a**), 15-30 nm (**b**), 30-45 nm (**c**), 45-60 nm (**d**), 60-75 nm (**e**) and 75-105 nm (**f**). Depth was measured from each particle center to the nearest milled surface.

**Supplementary Figure 7.**
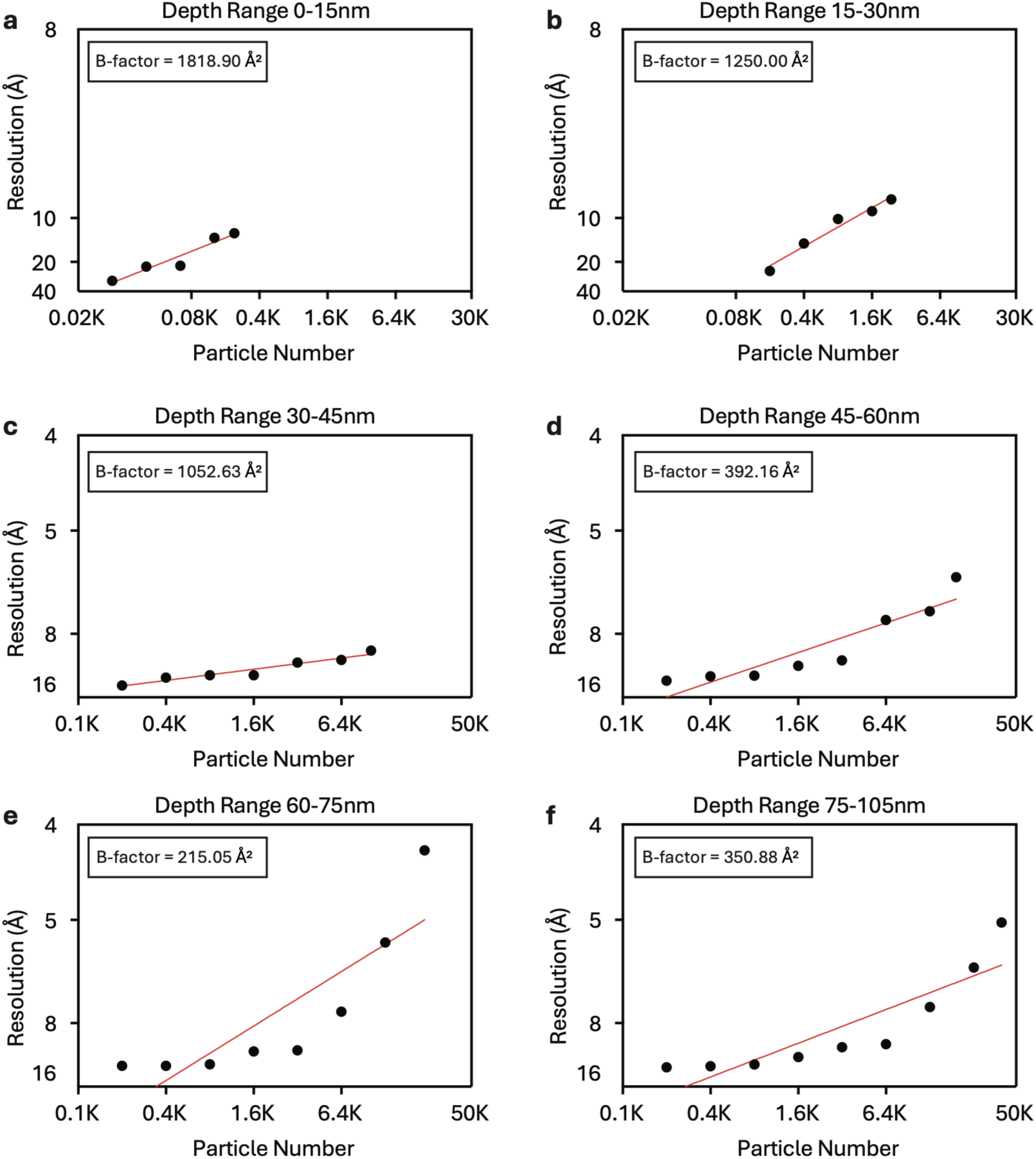
Rosenthal-Henderson analyses of apoferritin in argon-milled lamellae. a-f, Rosenthal-Henderson plots for *in situ* apoferritin from argon-milled *E. coli* lamellae, grouped by particle depth: 0-15 nm (**a**), 15-30 nm (**b**), 30-45 nm (**c**), 45-60 nm (**d**), 60-75 nm (**e**) and 75-105 nm (**f**). Depth was measured from each particle center to the nearest milled surface.

**Supplementary Figure 8.**
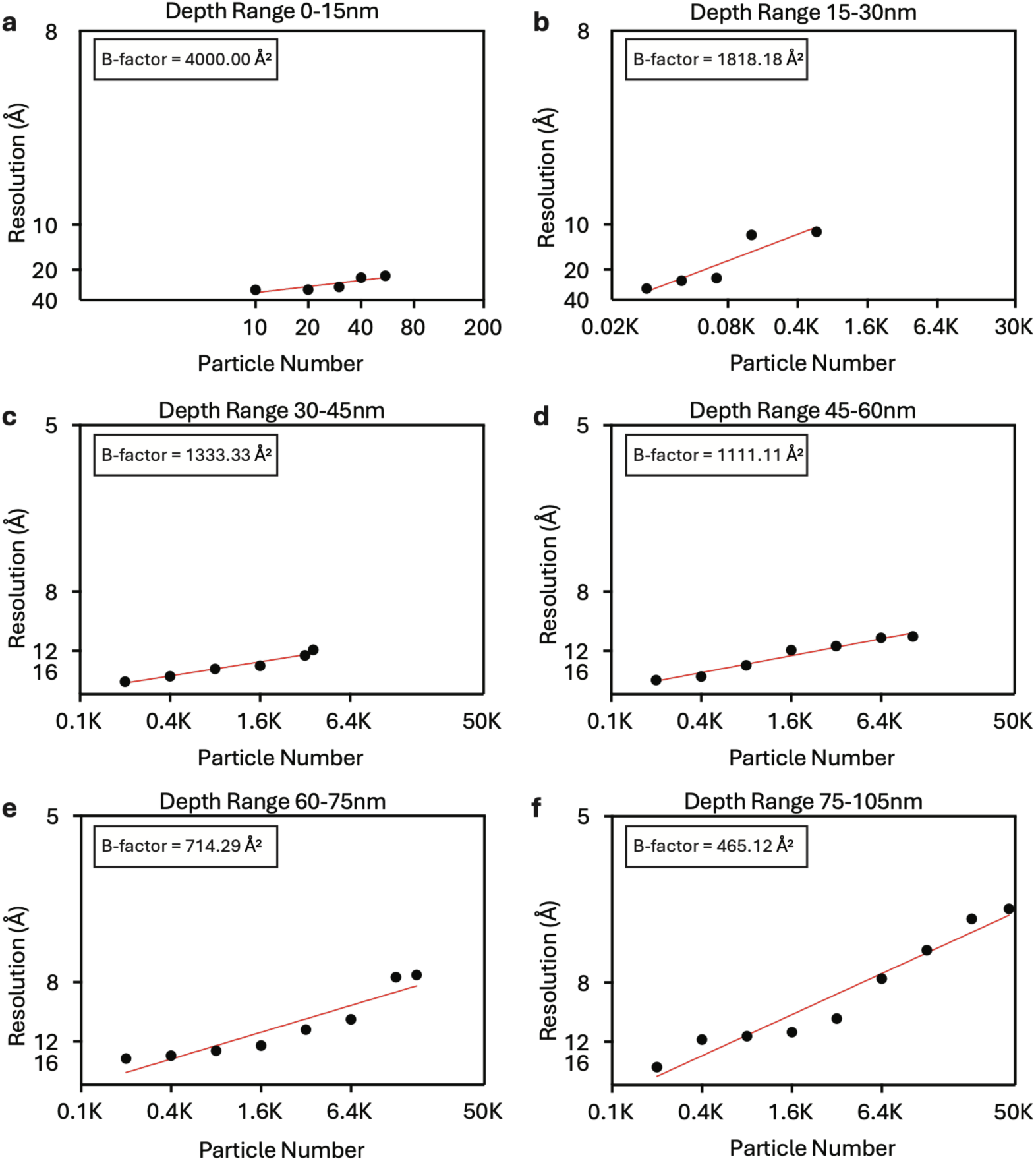
Rosenthal-Henderson analyses of apoferritin in oxygen-milled lamellae. a-f, Rosenthal–Henderson plots for *in situ* apoferritin from oxygen-milled *E. coli* lamellae, grouped by particle depth: 0-15 nm (**a**), 15-30 nm (**b**), 30-45 nm (**c**), 45-60 nm (**d**), 60-75 nm (**e**) and 75-105 nm (**f**). Depth was measured from each particle center to the nearest milled surface.

**Supplementary Figure 9.**
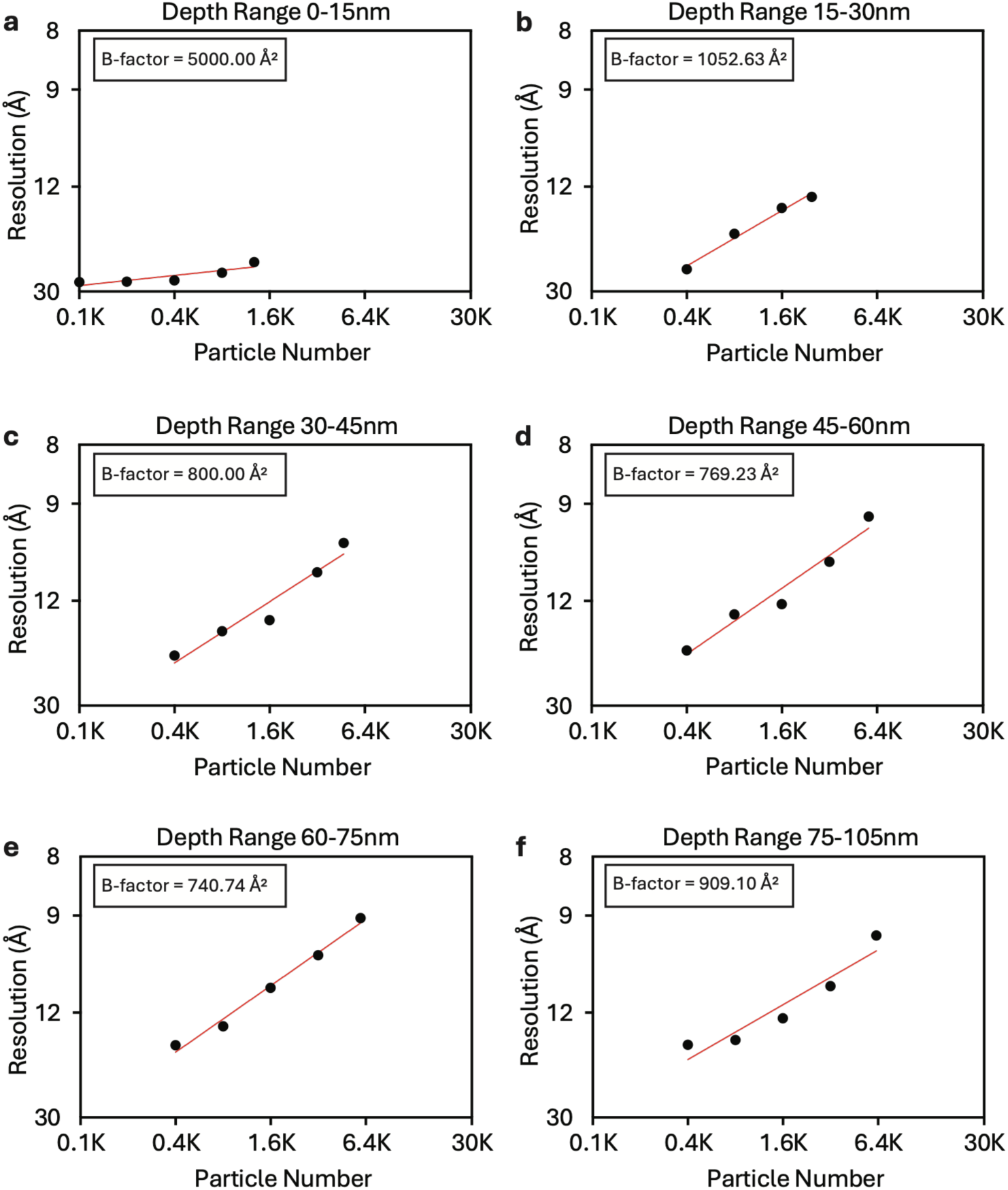
Rosenthal-Henderson analyses of β-galactosidase in xenon-milled lamellae. a-f, Rosenthal-Henderson plots for *in situ* β-galactosidase from xenon-milled *E. coli* lamellae, grouped by particle depth: 0-15 nm (**a**), 15-30 nm (**b**), 30-45 nm (**c**), 45-60 nm (**d**), 60-75 nm (**e**) and 75-105 nm (**f**). Depth was measured from each particle center to the nearest milled surface.

**Supplementary Figure 10.**
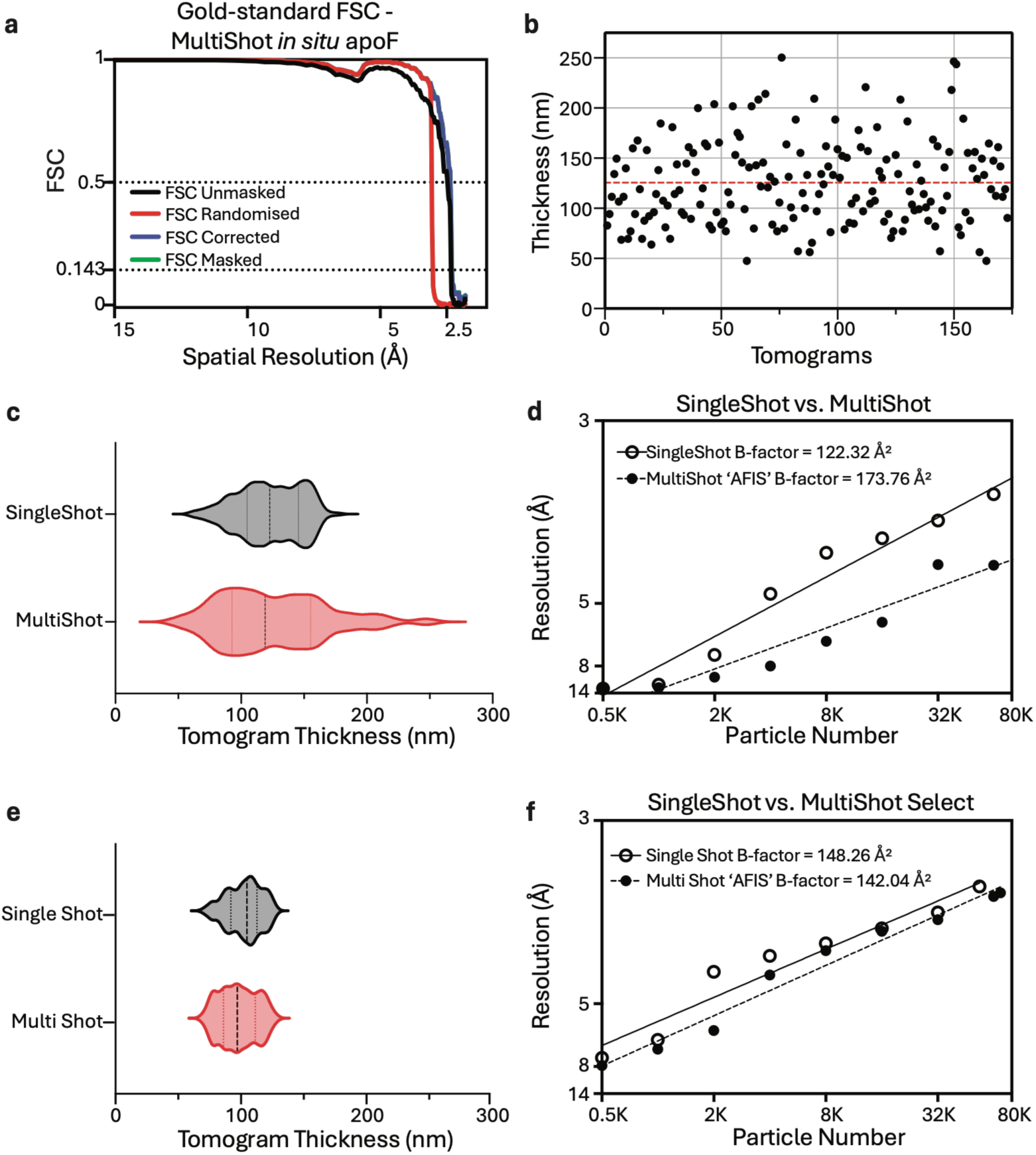
Comparison of single-shot and MultiShot acquisition for apoferritin using Tomography 5 software. a, Gold-standard Fourier shell correlation (FSC) curves for the MultiShot *in situ* apoferritin reconstruction, showing a resolution of 2.31 Å at FSC = 0.143. **b,** Estimated *E. coli* lamella thicknesses for the MultiShot dataset (125.5 ± 42.62 nm). The red dashed line indicates the mean. **c,** Lamella thickness distributions for the complete single-shot (gray) and MultiShot (red) datasets. **d,** Rosenthal-Henderson analyses comparing B-factors for the complete datasets. **e,** Lamella thickness distributions for subsets selected by restricting the thickness range. **f,** Corresponding Rosenthal-Henderson analyses for the selected subsets in **e**.

**Supplementary Figure 11.**
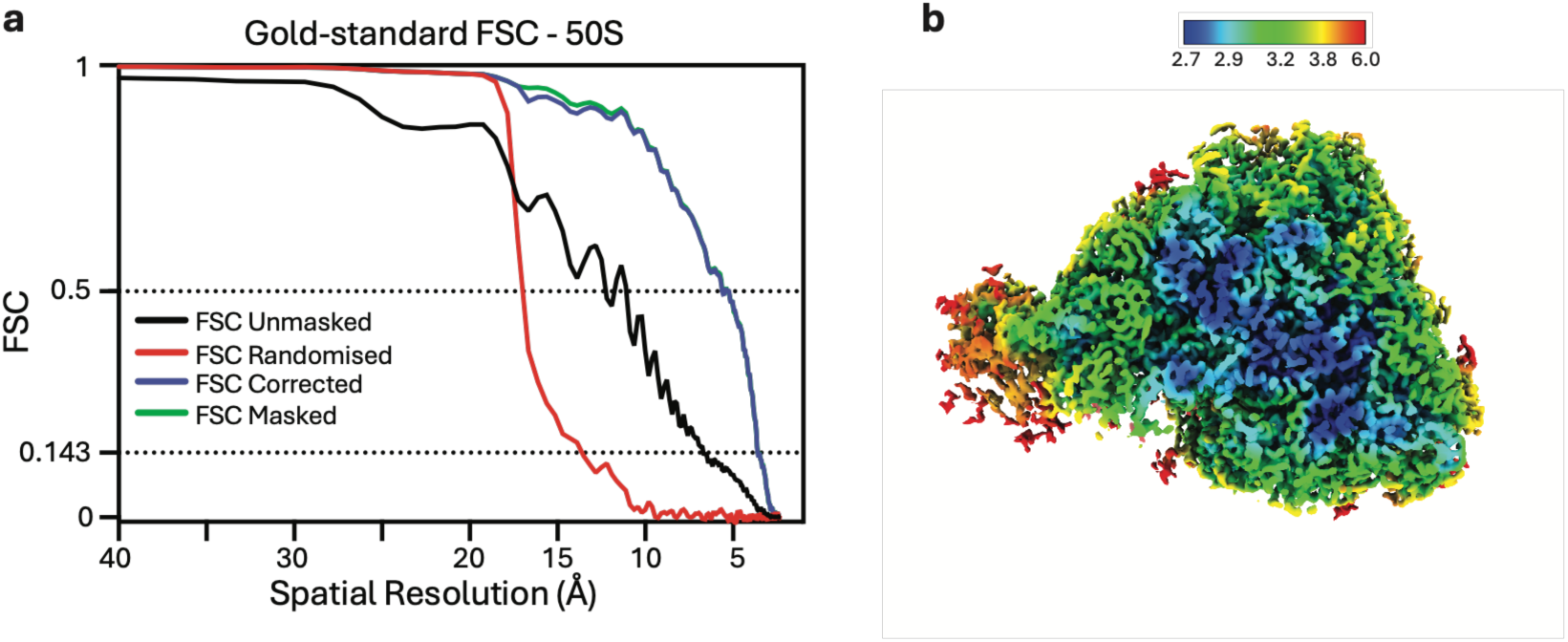
Global and local resolution of the 50S ribosomal subunit reconstruction. a, Gold-standard Fourier shell correlation (FSC) curves for the *in situ* 50S ribosomal subunit reconstructed from xenon-milled *E. coli* lamellae, showing a global resolution of 3.60 Å at FSC = 0.143. **b,** Corresponding reconstruction colored by local resolution (Å), with resolution better than 3 Å in parts of the map.

## Notes

### Competing Interest Statement

The authors have declared no competing interest.

## References

1. Nogales, E. and J. Mahamid, Bridging structural and cell biology with cryo-electron microscopy. Nature, 2024. 628(8006): p. 47–56.

2. McCafferty, C.L., et al., Integrating cellular electron microscopy with multimodal data to explore biology across space and time. Cell, 2024. 187(3): p. 563–584.

3. Dickerson, J.L., et al., Imaging biological macromolecules in thick specimens: The role of inelastic scattering in cryoEM. Ultramicroscopy, 2022. 237: p. 113510.

4. Rigort, A., et al., Focused ion beam micromachining of eukaryotic cells for cryoelectron tomography. Proceedings of the National Academy of Sciences, 2012. 109(12): p. 4449–4454.

5. Marko, M., et al., Focused-ion-beam thinning of frozen-hydrated biological specimens for cryo-electron microscopy. Nat Methods, 2007. 4(3): p. 215–7.

6. Schaffer, M., et al., A cryo-FIB lift-out technique enables molecular-resolution cryo-ET within native Caenorhabditis elegans tissue. Nat Methods, 2019. 16(8): p. 757–762.

7. Schaffer, M., et al., Cryo-focused Ion Beam Sample Preparation for Imaging Vitreous Cells by Cryo-electron Tomography. Bio-protocol, 2015. 5(17): p. e1575.

8. Berger, C., et al., Plasma FIB milling for the determination of structures in situ. Nature communications, 2023. 14(1): p. 629.

9. Berger, C., et al., Xenon plasma focused ion beam lamella fabrication on high-pressure frozen specimens for structural cell biology. Nature Communications, 2025. 16(1): p. 2286.

10. Kelley, R., et al., Toward community-driven visual proteomics with large-scale cryo-electron tomography of Chlamydomonas reinhardtii. Molecular Cell, 2026. 86(1): p. 213–230.e7.

11. Schiøtz, O.H., et al., Serial Lift-Out: sampling the molecular anatomy of whole organisms. Nature Methods, 2024. 21(9): p. 1684–1692.

12. Zens, B., et al., Lift-out cryo-FIBSEM and cryo-ET reveal the ultrastructural landscape of extracellular matrix. Journal of Cell Biology, 2024. 223(6): p. e202309125.

13. Creekmore, B.C., et al., Ultrastructure of human brain tissue vitrified from autopsy revealed by cryo-ET with cryo-plasma FIB milling. Nature communications, 2024. 15(1): p. 2660.

14. Lucas, B.A. and N. Grigorieff, Quantification of gallium cryo-FIB milling damage in biological lamellae. Proceedings of the National Academy of Sciences, 2023. 120(23): p. e2301852120.

15. Yang, Q., et al., The reduction of FIB damage on cryo-lamella by lowering energy of ion beam revealed by a quantitative analysis. Structure, 2023. 31(10): p. 1275–1281.e4.

16. Tuijtel, M.W., et al., Thinner is not always better: Optimizing cryo-lamellae for subtomogram averaging. Science Advances, 2024. 10(17): p. eadk6285.

17. Hall, L.A.-O.X., et al., Optimized cryo-FIB milling strategy to generate thin, minimally damaged biological lamellae. bioRxiv, 2026(2692-8205 (Electronic)).

18. Nakane, T., et al., Single-particle cryo-EM at atomic resolution. Nature, 2020. 587(7832): p. 152–156.

19. Tan, Y.Z. and B. Carragher, Seeing Atoms: Single-Particle Cryo-EM Breaks the Atomic Barrier. Molecular Cell, 2020. 80(6): p. 938–939.

20. Ali, M., et al., Characterization Standard for In-situ Cryo-electron Tomography. bioRxiv, 2026: p. 2026–05.

21. Guo, H., et al., Electron-event representation data enable efficient cryoEM file storage with full preservation of spatial and temporal resolution. IUCrJ, 2020. 7(Pt 5): p. 860–869.

22. Rosenthal, P.B. and R. Henderson, Optimal Determination of Particle Orientation, Absolute Hand, and Contrast Loss in Single-particle Electron Cryomicroscopy. Journal of Molecular Biology, 2003. 333(4): p. 721–745.

23. Kremer, J.R., D.N. Mastronarde, and J.R. McIntosh, Computer visualization of three-dimensional image data using IMOD. J Struct Biol, 1996. 116(1): p. 71–6.

24. Chaillet, M.L., et al. Extensive Angular Sampling Enables the Sensitive Localization of Macromolecules in Electron Tomograms. International Journal of Molecular Sciences, 2023. 24, 13375 DOI: 10.3390/ijms241713375.

25. Chaillet, M.L., et al., pytom-match-pick: A tophat-transform constraint for automated classification in template matching. Journal of Structural Biology: X, 2025. 11: p. 100125.

26. Bartesaghi, A., et al., Structure of β-galactosidase at 3.2-Å resolution obtained by cryo-electron microscopy. Proceedings of the National Academy of Sciences, 2014. 111(32): p. 11709–11714.

27. Bartesaghi, A., et al., 2.2 Å resolution cryo-EM structure of β-galactosidase in complex with a cell-permeant inhibitor. Science, 2015. 348(6239): p. 1147–1151.

28. Kimanius, D., et al., Accelerated cryo-EM structure determination with parallelisation using GPUs in RELION-2. eLife, 2016. 5: p. e18722.

29. Merk, A., et al., 1.8 A resolution structure of [beta]-galactosidase with a 200 kV CRYO ARM electron microscope. IUCrJ, 2020. 7(4): p. 639–643.

30. Danev, R., et al., Atomic resolution cryo-EM at 200 keV. IUCrJ, 2026. 13(4): p. 343–353.

31. Brogden, V., et al., Material Sputtering with a Multi-Ion Species Plasma Focused Ion Beam. Advances in Materials Science and Engineering, 2021. 2021(1): p. 8842777.

32. Dumoux, M.A.-O., et al., Cryo-plasma FIB/SEM volume imaging of biological specimens. 2023(2050-084X (Electronic)).

33. Totonjian, D.D., *Doped Beam Very Low Energy Particle Induced X-Ray Emissio*n. 2022, UTS Digital Thesis Collection.

34. Tegunov, D. and P. Cramer, Real-time cryo-electron microscopy data preprocessing with Warp. Nat Methods, 2019. 16(11): p. 1146–1152.

35. Zheng, S., et al., AreTomo: An integrated software package for automated marker-free, motion-corrected cryo-electron tomographic alignment and reconstruction. Journal of Structural Biology: X, 2022. 6: p. 100068.

36. Tegunov, D., et al., Multi-particle cryo-EM refinement with M visualizes ribosome-antibiotic complex at 3.5 Å in cells. 2021.

37. Goddard, T.D., et al., UCSF ChimeraX: Meeting modern challenges in visualization and analysis. Protein Science, 2018. 27(1): p. 14–25.

38. Watson, Z.L., et al., Atomistic simulations of the Escherichia coli ribosome provide selection criteria for translationally active substrates. Nature Chemistry, 2023. 15(7): p. 913–921.

39. Powell, B.M. and J.H. Davis, Learning structural heterogeneity from cryo-electron sub-tomograms with tomoDRGN. Nature Methods, 2024. 21(8): p. 1525–1536.

40. Burt, A., et al., An image processing pipeline for electron cryo-tomography in RELION-5. FEBS Open Bio, 2024. 14(11): p. 1788–1804.

41. Ermel, U.H., S.M. Arghittu, and A.S. Frangakis, ArtiaX: An electron tomography toolbox for the interactive handling of sub-tomograms in UCSF ChimeraX. Protein Science, 2022. 31(12): p. e4472.

